# Demographic determinants of biometric heritability

**DOI:** 10.1101/370866

**Authors:** Julia A. Barthold, Floriane Plard, Jean-Michel Gaillard, Tim Coulson, Shripad Tuljapurkar

## Abstract

The response of quantitative characters to selection depends on their transmission from parents to offspring. A common estimate of this transmission is the biometric heritability defined as the slope of the regression of offspring phenotype on same-aged mid-parent phenotype (i.e. the ratio of the phenotypic parent-offspring covariance over the parental phenotypic variance). This slope is often interpreted as the percentage of phenotypic variation due to additive genetic effects after accounting for confounding factors such as environment, litter or parental effects. However, researchers seldom account for the possible influence of selection on this estimate. Here we study the effect on biometric heritability of fertility and viability selection, as well as phenotype ontogeny (growth) and inheritance from parents to offspring. We present exact formulas for the elasticities of biometric heritability in age-phenotype-structured integral projection models (IPMs), and illustrate these for two iteroparous long-lived species. We find that both viability and fertility selection can strongly affect heritability, mediated by growth and inheritance. Generally, demographic processes that result in parents reproducing at large phenotypes, regardless of their own birth phenotype, decrease heritability. Analysed at equilibrium, our models imply that a heritable character can show no response to selection, if parental phenotypes affect offspring phenotypes and if phenotypes develop with age. Our results further highlight the importance of accounting for demographic processes when estimating heritability.

## Introduction

Evolutionary biologists seek to understand how traits evolve in natural systems. Predictions of trait evolution are derived from quantitative genetics theory (Falconer & Mackay, 1996). To evolve, a trait needs to vary, be under selection and transmitted to the next generation. A key measure of transmission, the parent-offspring phenotypic covariance, is usually interpreted as arising from genetic similarity between parents and their offspring. The ratio of parent-offspring phenotypic covariance over parental variance is the slope of the linear association between parental and offspring phenotypic traits measured at the same point in the life cycle. Once potential confounding factors affecting offspring phenotypic traits have been accounted for, this ratio is interpreted as the proportion of phenotypic variation attributable to additive genetic variation—the narrow-sense heritability, or *h*^2^, of a trait (Falconer & Mackay, 1996; Kempthorne, 1957). The reason underpinning this logic is that genotypes are inherited at birth and remain constant throughout life, with the offspring genotype expected to be intermediate between the genotypes of the two parents (Lynch & Walsh, 1998). Heritability enters the breeder’s equation to predict shifts in the population mean of a quantitative character under selection from one generation to the next. However, this interpretation of the parent-offspring covariance is restrictive. It excludes non-genetic inheritance (Bonduriansky & Day, 2009; Danchin et al., 2011) and ignores trait ontogeny. However, individuals must grow and be sexually mature before reproducing and the resulting parental conditions influence offspring phenotype (Mousseau & Fox 1998, Figure 1).

**Figure 1:**
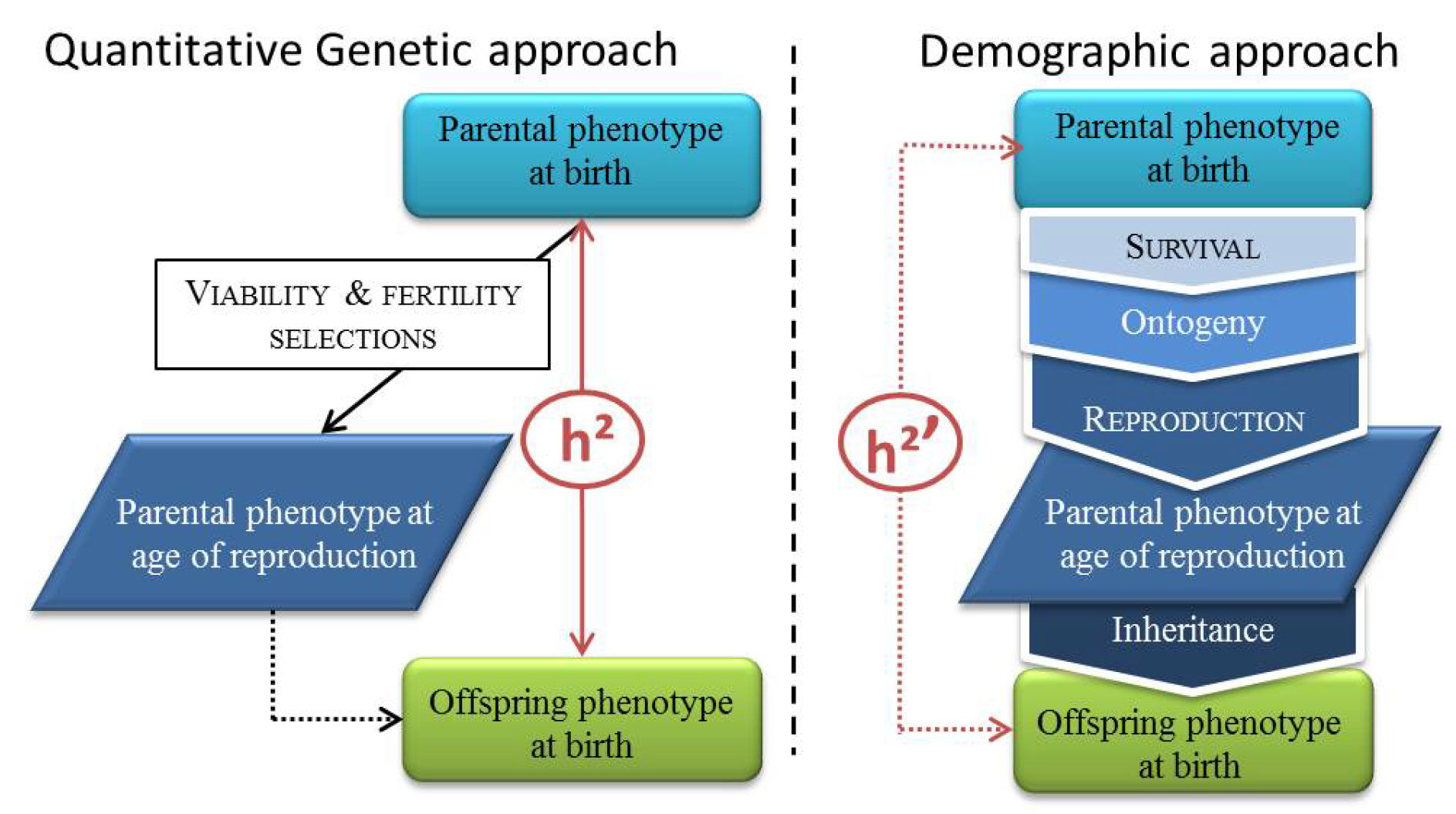
Differences between heritabilities directly measured from data and parent-offspring regression (*h*^2^) and derived from IPMs (*h*^2^’). In the demographic approach, survival and ontogeny covary during the growing stage.

A key observation challenges the widespread use of the breeder’s equation to study evolution in the wild. Size-related traits are frequently heritable and under directional selection in free-living populations of animals and plants, but they do not evolve as predicted by the breeder’s equation (Merilä et al., 2001). The reason for this is that many assumptions underlying this equation are likely violated in natural systems. For instance, incomplete or flawed pedigrees can bias the estimation of breeding values (Postma, 2006). Furthermore, the component linked to the interaction between genes and environment cannot be estimated in empirical studies in the wild and is therefore often assumed to be negligible. Finally, the link between phenotype and fitness can be caused by a correlated but unknown environmental or phenotypic trait (Morrissey et al., 2010).

In contrast to the challenges met when applying the breeder’s equation to predict trait development, eco-evolutionary demographers have developed data-driven population models (Integral projection models, IPM,Easterling et al. 2000) that match observed phenotypic change over time (Coulson et al., 2011; Smallegange & Coulson, 2012; Vindenes et al., 2014). However, these models do not comply with quantitative genetics theory (Chevin 2015, but see Coulson et al. 2017), they do not follow the genotype and do not assume that the phenotype is influenced solely by additive genetic and environmental effects. IPMs follow the phenotype distribution and its dynamics in populations. They include a purely phenotypic across-age notion of inheritance (Janeiro et al., 2016). In this way, IPMs capture maternal and environmental effects of phenotype transmission through an inheritance function that links phenotype of the mother at reproduction to the phenotype of the offspring (see Fig. 1). Moreover, this approach explicitly includes ontogeny so that a new-born first grows to reach a required size at maturity, then reproduces, and lastly transmits its phenotype to its offspring. Coulson et al. (2017) have recently demonstrated how IPMs can track the additive genetic and environmental component of the phenotype, similar to approaches used in quantitative genetic models. But here, we suggest studying how ontogeny and selection can influence the measure of heritability using IPMs tracking directly the phenotype dynamics.

As long as major challenges exist for the application of quantitative genetics theory to natural systems, and without discrediting one or the other approach, the aim of this paper is to study heritability using these data-driven population models in order to stay open to the question how other processes than genetic inheritance could affect estimates of heritability. We calculate heritability from the modelled phenotype distributions at equilibrium as the ratio of parent-offspring phenotypic covariance over parental variance, which is the exact same mathematical definition of the slope of the parent-offspring regression as developed in quantitative genetics theory (Falconer & Mackay, 1996). While the estimate of the slope is usually corrected for non-genetic effects, when calculated from an IMP it explicitly includes both genetic and non-genetic mechanisms of inheritance. IPMs furthermore explicitly model ontogeny to estimate the parent-offspring covariance. To indicate that heritabilities reported here contain non-genetic mechanisms of inheritance, and are affected by fertility and viability selection and growth, we denote the quantity as “biometric heritability” (after Jacquard, 1983).

To study biometric heritability, we begin with a general IPM that describes changes in a trait (e.g. body mass) with age. For any such IPM, we derive analytical expressions for biometric heritability and for the elasticity of heritability to changes in any model parameter (intercept and slope of the different functions constituting the IPM). Elasticity of heritability measures the proportional change in heritability caused by an infinitesimal change of one of the model parameter and is estimated using the derivative of heritability with respect to this trait over heritability. These expressions will apply to any IPM of this type, and can be extended to IPMs that describe multivariate traits. We apply our general results to analyse two published age-body mass-structured IPMs, one parameterised for Soay Sheep (*Ovis aries*) (Coulson et al., 2010) and one for roe deer (*Capreolus capreolus*) (Plard et al., 2015). In these iteroparous species, body mass is under positive selection and develops with age. Surviving parents produce offspring that inherit a birth phenotype, which may be a function of the environment and parental age, phenotype, and condition (Mousseau & Fox, 1998; Skibiel et al., 2009). As a consequence, the parent-offspring phenotypic covariance arises out of complex interactions between ontogeny, demography, and selection. We report and compare how the biometric heritability of body mass for both species depends on selection processes, growth, and inheritance. By doing this, we show how demographic processes could influence estimates of heritability.

## Methods

The **biometric heritability** *h*^2^ of a phenotypic trait is the slope of the regression of offspring phenotype values *Y_a_* (“offspring phenotype”) at age *a* on mid-parent phenotype values *X_a_* (“parent phenotype”) at age *a* (Jacquard, 1983) (referred to as “heritability” from now on). We estimate heritability of traits at birth using phenotypic and life history information from all ages. Our approach can be used for traits measured at any age, which have to be the same for parents and offspring. Coulson et al. (2010) generate parent-offspring phenotype patterns using IPMs and estimate the regression slope of the phenotypes of offspring born to one parent cohort over its lifetime regressed on the phenotypes of parents at their own birth. The heritability can be calculated using the formula for a regression slope

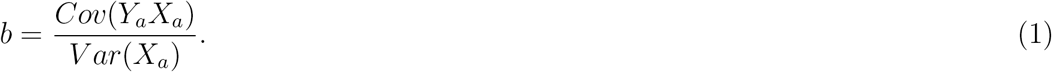

If parents of only one sex are considered, the heritability *h*^2^ equals twice the slope *b*, since one parent only contributes half of the offspring genome (Falconer & Mackay, 1996). From (1), it follows that the **elasticity** of the biometric heritability, defined as the proportional change in *b*, is

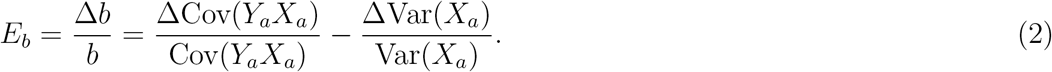

The **elasticity** of heritability *E_b_* is the derivative of the heritability ∆*b* with respect to one of the parameter of the model, divided by this heritability. The precise forms of the elasticity of heritability *E_b_* depend on which model parameter is changed. The quantities in (1) and (2) can be derived for an age-phenotype-structured population model, where phenotype is a quantitative character whose phenotypic values are discrete elements of the vector **z** = {*z_i_*}. In the following, we describe how the quantities for calculating the elasticity of heritability can be computed. Detailed derivations of the described quantities are given in the supplemental material (equations A1-A9).

### Stable cohort birth phenotype distribution

The age and phenotype structure of a population that experiences time-invariant fertility and mortality rates converges to a stable age and phenotype distribution with a growth rate *r* (Keyfitz & Caswell, 2005). Stable populations have therefore stable proportions of newborns in each phenotype class, here represented by a vector **u** = {*u*(*i*)} ≡ {*u*(*z_i_*)} and referred to as the **stable cohort birth phenotype distribution**, which contains all the newborns of a given year. Note that we set (**e**^*T*^ **u**) = 1, using the vector **e**^*T*^ = (1, 1*,…,* 1). If we define for any *r* a renewal matrix

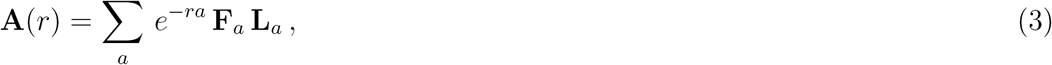

where **F**_*a*_ is a matrix describing fertility for parents of age *a*, and **L**_*a*_ is a matrix describing survivorship from birth to age *a* (both matrices are described in detail in the next section), then the stable cohort birth phenotype distribution **u** and growth rate *r* are together determined by

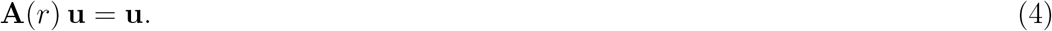

For details on the renewal matrix and the stable cohort birth phenotype distributions see Steiner et al. (2014).

### Phenotype-demography matrices

**Fertility** of parents aged *a* is a matrix 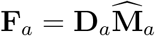, where 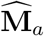 is a diagonal matrix whose (*i, i*) entry is the total number of offspring that a parent of phenotype *z_i_* will produce at age *a*. The **inheritance** of the phenotype from a parent of phenotype *z_i_* at age *a* to offspring is captured in the parent-offspring phenotypic transition matrix **D**_*a*_ whose (*i, j*) element is

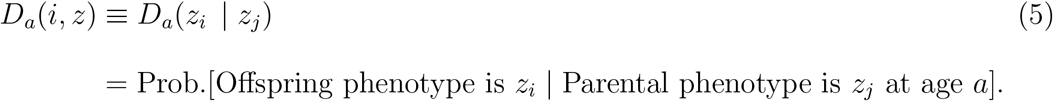

Note that **e**^*T*^ **D**_*a*_ = **e**^*T*^ so that the probabilities for the transitions out of one parental phenotype *z_j_* sum to one.

**Survival and growth** together from age *a* to age *a* + 1 are given by a matrix **P**_*a*_ whose (*i, j*) element is

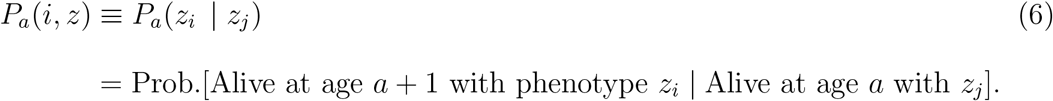

**Survivorship** from birth at age *a* = 0 to any age *a* is a matrix **L**_*a*_ whose (*i, j*) element is

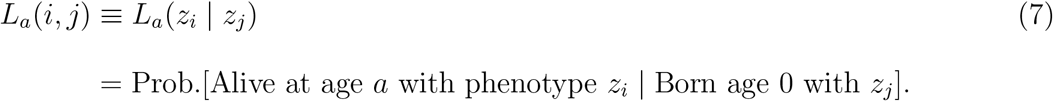

### Parent-offspring covariance and parental variance

In the first part of the appendix (A1) we derive a formula for the covariance between offspring phenotype *Y* and parent birth phenotype *X* that is computable in terms of **F**_*a,*_ **L**_*a*_, and **u**. As a consequence, the parent-offspring phenotypic covariance arises out of complex interactions between ontogeny, demography, and selection.

In a stable age-phenotype-structured population, the **lifetime number of offspring** produced by a birth cohort that has the stable cohort birth phenotype distribution **u** is

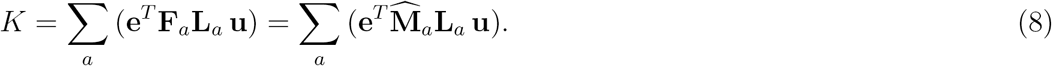

Using this, the phenotype-demography matrices **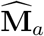** and **L**_*a*_, and the stable cohort birth phenotype distribution **u** as defined in (4), the **parent-offspring covariance** between offspring and parent birth phenotypes is given by

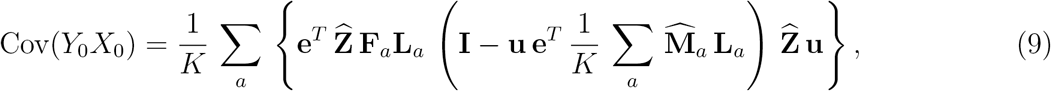

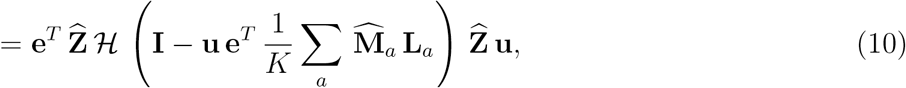

where **Z**̂ is a diagonal matrix of the phenotypic values **Z**̂ = *diag*(*z_i_*), and where we use, for brevity,

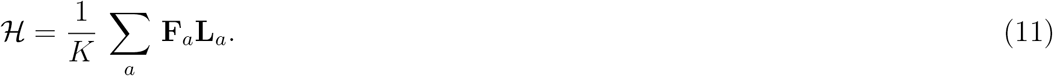

Details on the derivations are given in the appendix A1, particularly in (A-14 to A-17). **The variance in parent birth phenotype** is

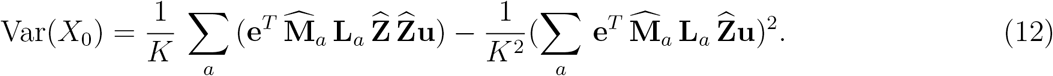

Details on the derivation of the variance is given in the appendix A2, particularly in (A-18 to A-23).

### Computing the elasticity of heritability

A change in biometric heritability ∆*b* occurs if we change the model parameters that govern the strength of the phenotype selection and transition processes by a small amount ∊ *>* 0. This change was estimated by the derivative of *b* in relation to the parameter that has changed. This causes changes in the phenotype-demography matrices

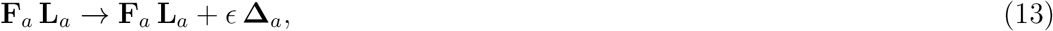

where **∆** represent the derivative of a matrix. In the following, we also use ∆ and ***δ*** to represent the derivative of a number and a vector, respectively. The appendix A4 and A9 give the details on the computation of the changes in the phenotype-demography matrices with respect to changes in viability selection, fertility selection, growth, and inheritance.

The changes in (13) change the stable population so that the stable cohort birth phenotype distribution **u** changes by ∊***δ***_*u*_ (see A3, A6, A7 and A8 for the computation of ***δ***_*u*_). From here on, we leave off the ∊ (constant proportional factor) since it multiplies every change. In addition to changes in **u**, the parent-offspring covariance Cov(*Y*_0_*X*_0_) changes by ∆Cov(*Y*_0_*X*_0_) and the variance in parent birth phenotype Var(*X*_0_) by ∆Var(*X*_0_) (see A5 for details on the perturbations of the parent birth phenotype variance).

The perturbation of the parent-offspring covariance as given in (10) requires the perturbation of matrix ℋ (11), which in turn needs the perturbation of *K* (8). From (11) see that the change in ℋ is

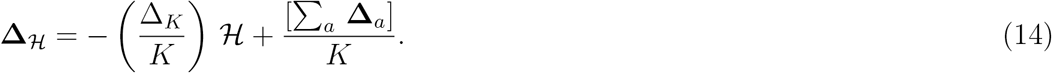

And from (8) we have

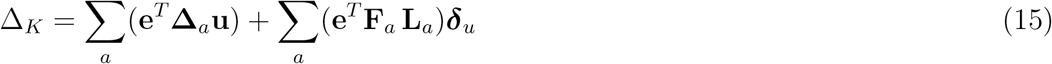

Putting these together, the change in parent-offspring covariance between offspring and parent birth phenotypes using (10) and ***δ***_*u*_ (A3) is

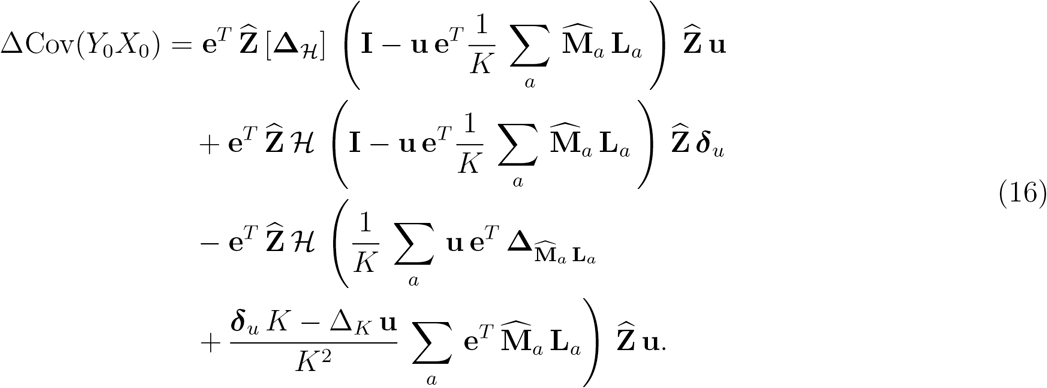

Finally, the change in parent birth phenotypic variance (Appendix A5) is

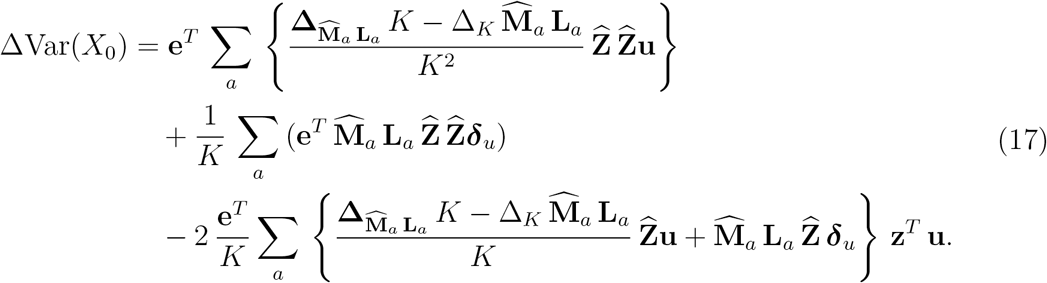

Using (10), (12), (16), and (17), we can now compute the biometric heritability in (1) and its elasticity in (2).

We also compare our analytical framework to numerical evaluation of elasticities of heritability with respect to changes in the model parameters. We numerically computed the elasticities from simulations by adding 10^*−*5^ to each model parameter, one at a time, dividing the resulting change in the perturbed elasticity by the unperturbed elasticity, and then scaling it back up by 10^5^. This approach is described in detail in Coulson et al. (2010).

We analyse the model at its equilibrium state when the phenotype distribution has stable, time-invariant proportions in each age and phenotype class. This stable phenotype distribution changes if model parameters are altered. As a consequence, changes in model parameters can affect heritability at population equilibrium by affecting one of three phenotype distributions: the stable cohort at birth phenotype distribution, the stable parent (at own) birth phenotype distribution, or the offspring birth phenotype distribution. The offspring produced by a stable population in one time step has the stable cohort birth phenotype distribution. Each such cohort contains individuals who may become parents if they survive and reproduce. Which of these potential parents reproduce, and how often, during their lifetimes determines the stable parent (at own) birth phenotype distribution. The latter is shifted to higher phenotype values when compared to the stable cohort birth phenotype distribution, if viability and fertility selection are positive. Finally, the offspring born to one cohort of parents over their lifetimes form the offspring birth phenotype distribution.

### Application to Soay sheep and roe deer IPMs

We computed the biometric heritability of body mass in Soay sheep and roe deer and the elasticities of heritability to changes in model parameters for two published age-phenotype-structured IPMs (Coulson et al., 2010; Plard et al., 2015). Both IPMs used data from the female component of the population and categorised those ages with statistically indistinguishable fertility, survival, growth, and inheritance functions into age-classes. The Soay sheep IPM divided individuals into four age-classes: lambs (aged 0 to *<* 1, census at 4 months old), yearlings (aged 1 to *<* 2), adults (aged 2 to *<* 8), and senescent individuals (aged 8+). Whereas the roe deer IPM distinguished three age-classes: yearlings (aged 0 to *<* 1, census at 8 month old), adults (aged 1 to *<* 7), and senescent individuals (aged 7+). For consistency among the IPMs and the above notation, where age 0 refers to age at birth (i.e. census age), we define the roe deer age-classes starting at age 0 to denote the census age of 8 months, instead of age 1 as in Plard et al. (2015). This is purely a question of notation. For both species, we calculate heritability for their respective census ages of 4 months in Soay sheep and 8 months in roe deer.

The IPM phenotype-demography matrices were constructed by predicting from the relevant phenotype-demography functions. The functions were statistically determined using life history and body mass data while controlling for confounding temporal variation in demographic rates. The functions inferred, by body weight and age-class, were yearly probability of survival, yearly probability of recruitment, probability of twinning, mean and variance of annual growth, and mean and variance of parent-offspring body weight inheritance. The survival, probability of recruitment, and twinning rate functions were estimated with a logit transformation while the mean and the variance of the growth and the inheritance functions used linear functions. Every function has an intercept and a slope per age-class. The total number of model parameters is the total number of coefficients of all functions. For a detailed description of the construction of IPMs see Merow et al. (2014) or Rees et al. (2014).

We now provide a brief summary of the data. The feral population of Soay sheep lives on the Island of Hirta in the St. Kilda archipelago, Scotland, and fluctuates as a food-limited population between 600 and 2000 individuals, of which about one-third live in the 250ha study area (Clutton-Brock & Pemberton, 2004). The population has been studied in detail since 1985. During this time, life-history and body mass data were collected during the yearly capture and year-round censuses. A detailed description of the population and data collection protocols is provided elsewhere (Clutton-Brock & Pemberton, 2004).

The roe deer population of Trois Fontaines lives in an enclosed forest of 1360 ha in North Eastern France (48°3*’*N, 2*°*61*’*W). The size of the population has been kept relatively constant around 250 individuals by yearly removals (mainly through exportation of captured individuals) except between 2001-2005 when an experimental manipulation of density was performed and population size peaked at 450 individuals. The population has been monitored by the Office National de la Chasse et de la Faune Sauvage since 1975. Each year, half of the population of roe deer is caught between December and March. Each individual is sexed and weighed. All individuals included in this study are of known age. The population and the study site has been described in details by Gaillard et al. (1993, 2013).

## Results

The estimated heritabilities of body mass were 0.20 for Soay sheep and 0.34 for roe deer. While the parent birth phenotypic variances (denominator used in the estimation of heritability) were similar (5.59 and 5.39 for roe deer and Soay sheep, respectively), the parent-offspring covariance (numerator) was smaller in Soay sheep (0.53) than in roe deer (0.94). This is because Soay sheep can give birth at 1 year of age and so give birth to offspring of different body masses at different ages, whereas roe deer give birth from 2 years of age onwards to more similar offspring each year. The elasticities of the heritabilities varied between −6% and 2.5%. Both the analytically and numerically calculated elasticities provided equivalent results (Figs. S1 and S2). In general, perturbing slopes had larger effects on heritability than perturbing intercepts (Figs. 2 and 3). In the following, we will discuss the analytically calculated elasticities with respect to changes in fertility and viability selection, growth, and inheritance. To understand the elasticities of heritability, we also discuss the underlying elasticities in the means and variances of the three phenotype distributions that contribute to determining heritability: the stable cohort birth phenotype distribution, the parent birth phenotype distribution, and the offspring birth phenotype distribution (Tables 1, S1, and S2).

**Table 1:** Change in key population parameters (%): expected and variance values of the stable cohort phenotype distribution (*X_c_*_0_), the stable parent (at own) birth phenotype distribution (*X*_0_), and offspring birth phenotype distribution (*Y*_0_) when selected model parameters are perturbed (see Tables S1 and S2 for perturbations of all parameters). Note that shown values were calculated numerically by adding 0.01 to model parameters.

| | | $\mathcal{E}(X_{c_0})$ | $\text{Var}(X_{c_0})$ | $\mathcal{E}(X_0)$ | $\text{Var}(X_0)$ | $\mathcal{E}(Y_0)$ | $\text{Var}(Y_0)$ | $\text{Cov}(Y_0X_0)$ | $h^2$ | Effect<sup>†</sup> |
| --- | --- | --- | --- | --- | --- | --- | --- | --- | --- | --- |
| Soay sheep | SurvSlpL* | 0.00 | -0.30 | -0.15 | -0.71 | 0.15 | -1.24 | -4.38 | -3.69 | -- |
|  | SurvSlpY | 0.01 | -0.09 | -0.07 | 0.05 | 0.05 | -0.38 | -1.73 | -1.79 | - |
|  | SurvSlpA | 0.04 | -0.14 | -0.01 | -0.11 | 0.15 | -0.94 | -3.75 | -3.65 | -- |
|  | PrSlpL | -0.41 | 2.84 | -0.13 | 1.37 | -0.35 | 2.59 | 4.53 | 3.12 | ++ |
|  | PrSlpA | 0.15 | -0.03 | 0.14 | 0.11 | 0.40 | -1.58 | -4.11 | -4.21 | -- |
|  | GrMSlpA | 0.30 | 1.55 | 0.39 | 2.01 | 0.38 | 0.90 | -0.71 | -2.67 | -- |
|  | GrMSlpS | -0.04 | -0.15 | -0.03 | -0.14 | -0.01 | -0.57 | -2.09 | -1.95 | - |
|  | InMSlpY | 0.13 | -0.10 | 0.09 | -0.29 | 0.12 | -0.17 | 1.45 | 1.75 | + |
|  | InMSlpA | 1.30 | 2.55 | 1.25 | 2.13 | 1.35 | 1.87 | 1.52 | -0.60 | - |
| roe deer | SurvSlpY | -0.11 | 0.00 | -0.17 | -0.12 | -0.03 | 0.07 | -0.09 | 0.02 | 0 |
|  | SurvSlpA | 0.13 | -0.13 | 0.02 | -0.16 | 0.36 | -0.10 | -6.02 | -5.87 | -- |
|  | PrSlpA | -0.27 | -0.14 | -0.29 | -0.15 | -0.16 | -0.03 | 0.86 | 1.01 | + |
|  | GrMSlpY | 0.32 | -0.68 | 0.28 | -0.69 | 0.30 | -0.72 | 0.50 | 1.21 | + |
|  | InMSlpA | 1.64 | -0.71 | 1.51 | -0.84 | 1.54 | -1.33 | 0.70 | 1.56 | + |

\* Model parameter labels are written in CamelCase forming compounds of the following abbreviations in order of appearance: survival (Surv), slope (Slp), lambs (L), yearlings (Y), adults (A), probability of recruitment (Pr), growth mean (GrM), senescents (S), inheritance mean (InM)). <sup>†</sup>Effect size classification based on the elasticities of $h^2$ : minimum to -2 (--), -2 to 0.25 (-), 0.25 to +0.25 (0), 0.25 to 2 (+), 2 to maximum (++).

**Figure 2:**
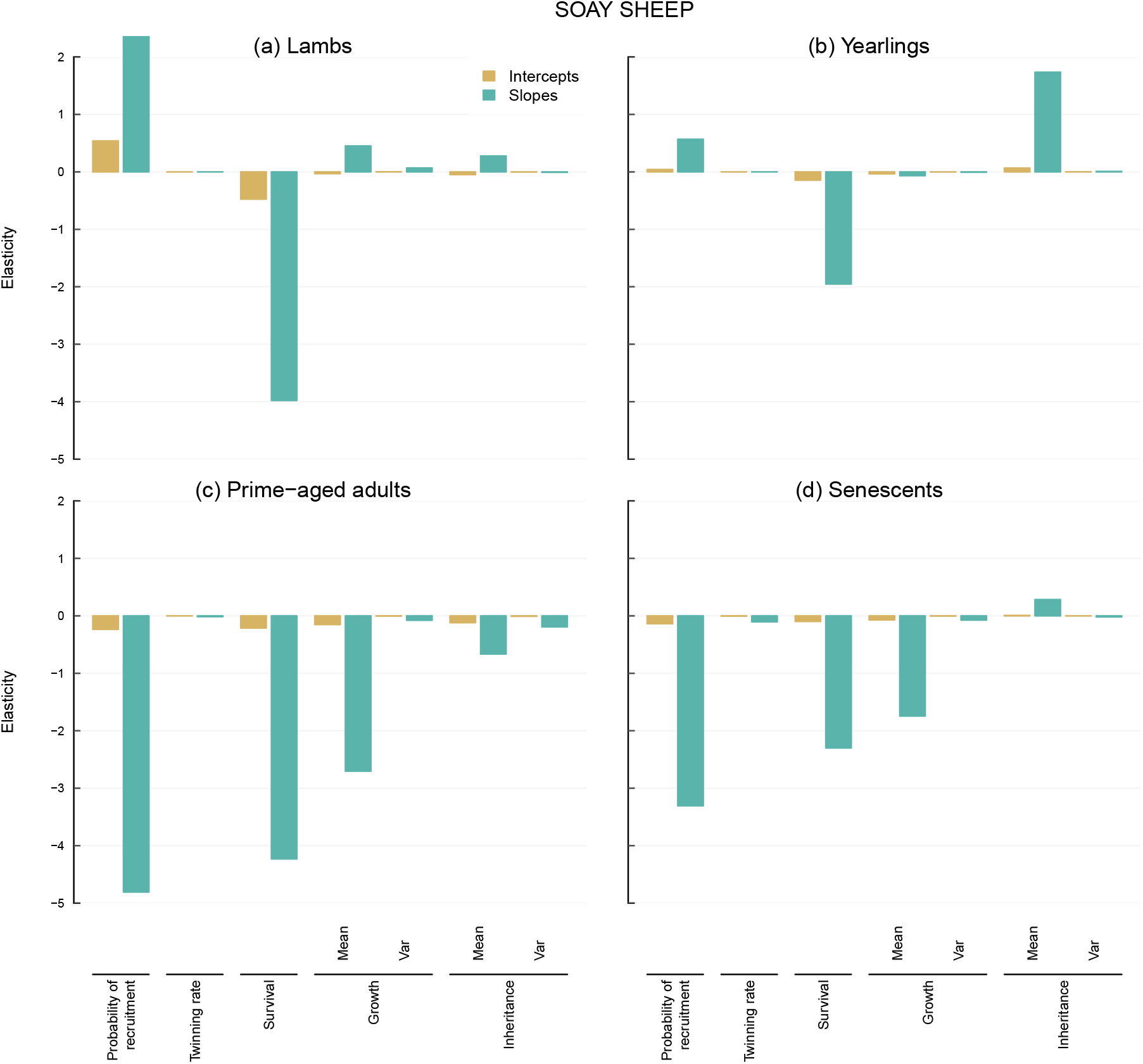
The analytically derived elasticities of heritability with respect to changes in the model parameters of the Soay sheep IPM.

**Figure 3:**
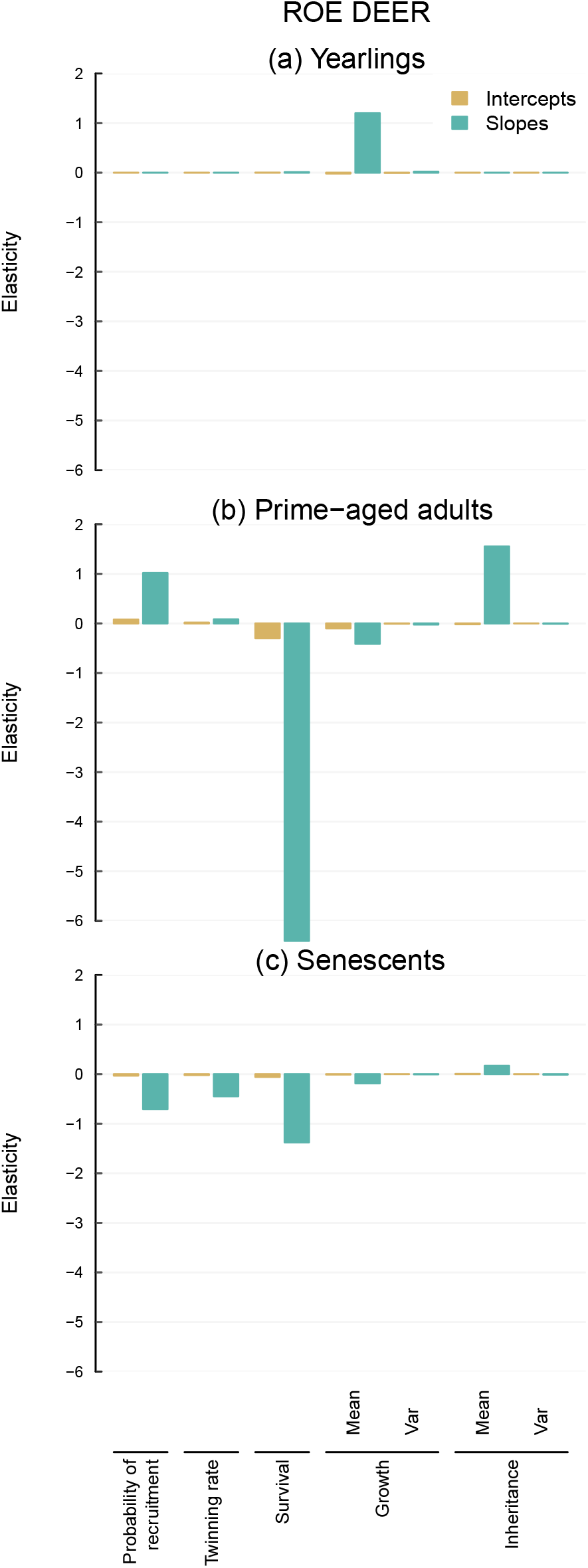
The analytically derived elasticities of heritability with respect to changes in the model parameters of the roe deer IPM.

### Elasticities of heritability with respect to viability selection

Increasing the slopes of the survival function and thus increasing survival and altering viability selection resulted in large negative elasticities of heritability for both species and almost all age-classes (Figs. 2 and 3). Changes to the slopes for the adult age-classes had the most pronounced effects. Increasing these slopes increased survival across all phenotype values but decreased the strength of viability selection because these changes increased survival of small adults more than survival of large adults. This is because the survival probabilities predicted by the unperturbed logit survival models were already approaching the boundary of one for larger adults (Figs. 4, 5, also see Fig. S3 for a schematic of this “linearisation effect”). In contrast to this, increasing the survival slope for Soay sheep lambs, for example, increased survival and the strength of viability selection, meaning that survival increased more for larger lambs than for smaller lambs (Figs. S3 and 4).

**Figure 4:**
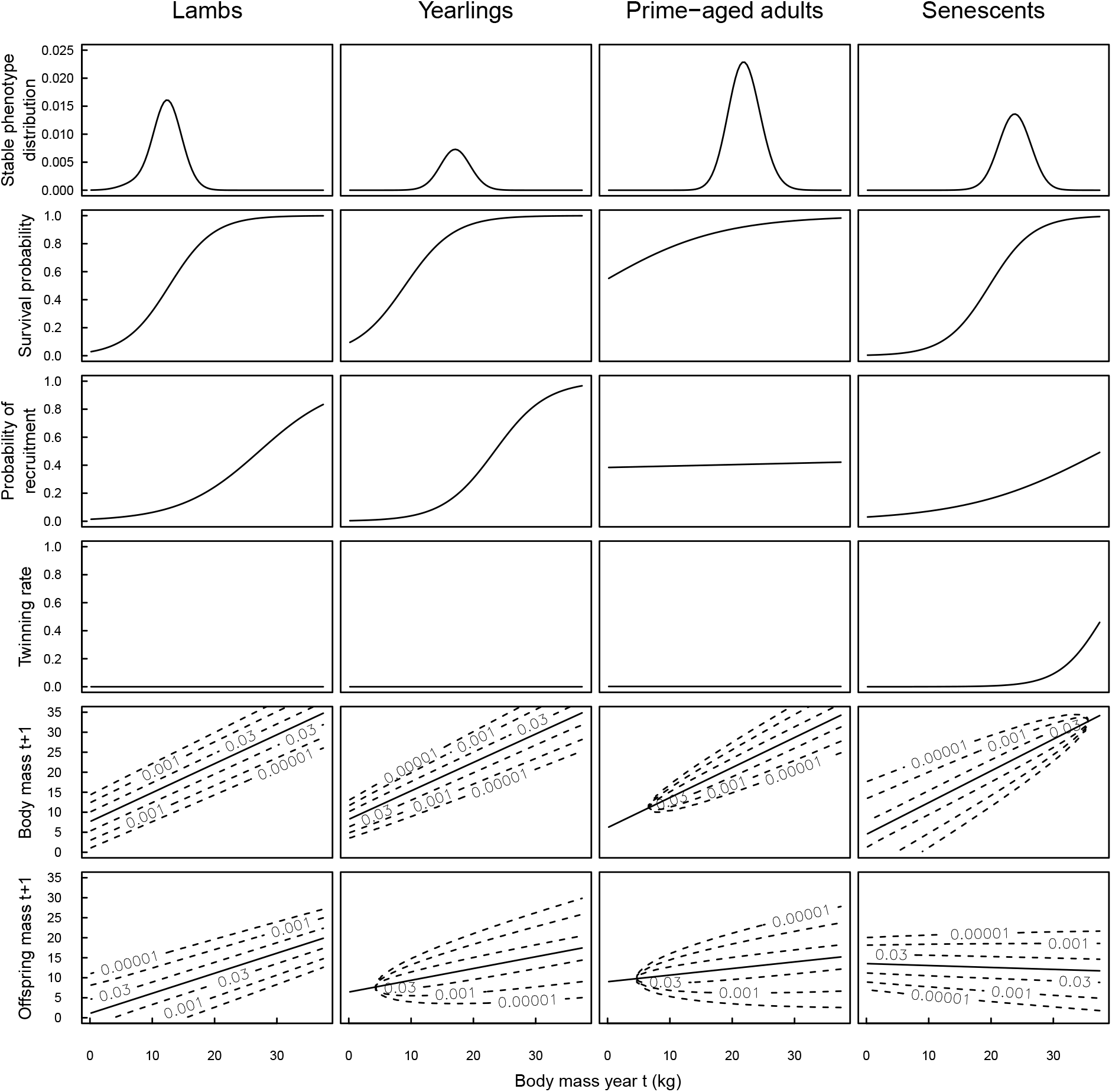
The phenotype-demography relationships used to parameterise the matrices in the IPM and the predicted stable phenotype distributions for Soay sheep. The age-class phenotype distributions together sum up to 1. For the phenotype-demography functions, lines represent predictions from regressions and dashed contours distributions around the mean.

**Figure 5:**
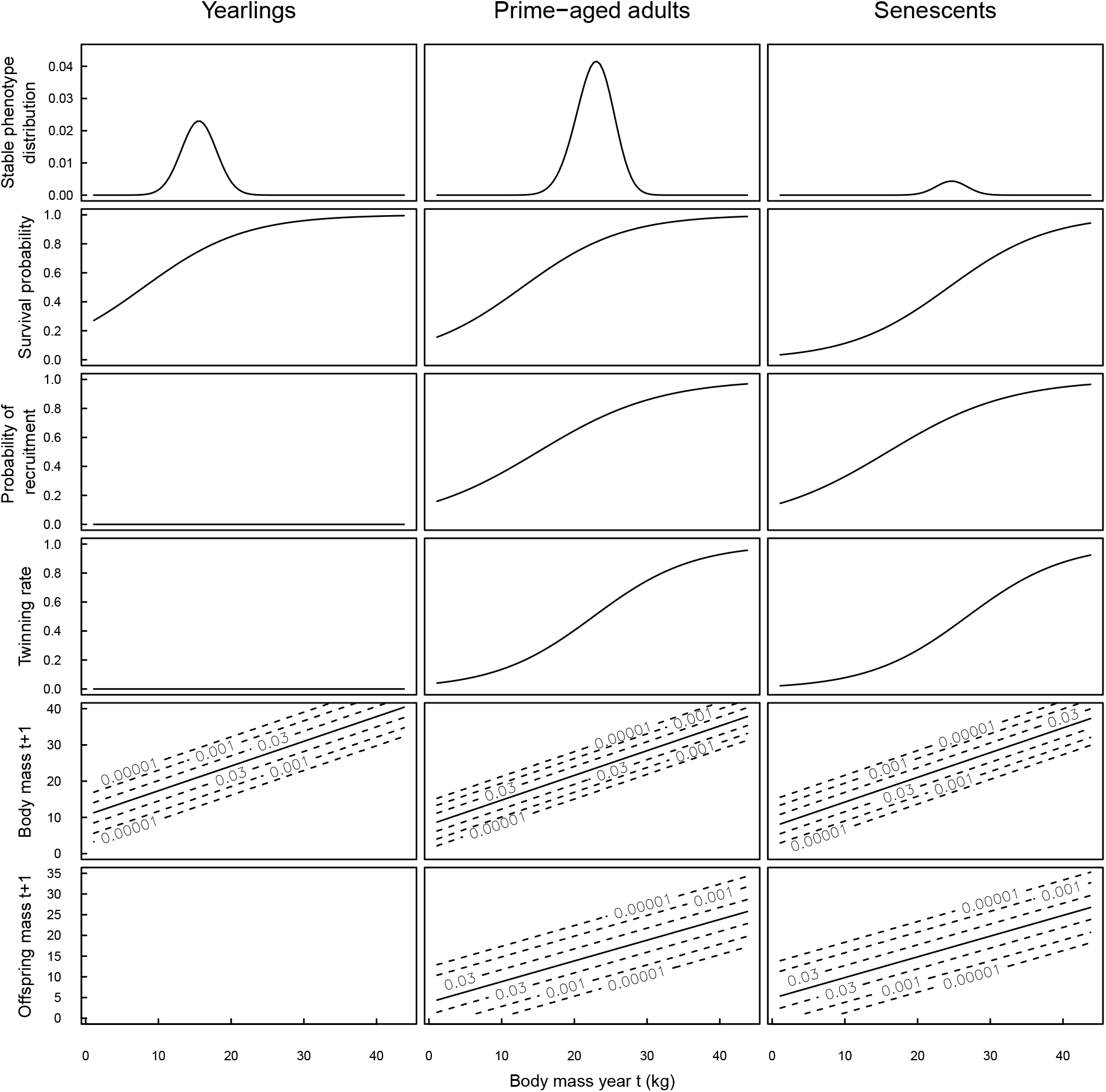
The phenotype-demography relationships used to parameterise the matrices in the IPM and the predicted stable phenotype distributions for roe deer. The age-class phenotype distributions together sum up to 1. For the phenotype-demography functions, lines represent predictions from regressions and dashed contours distributions around the mean. Roe deer yearlings do not reproduce.

Regardless of the direction of the change in viability selection, increasing the adult survival slopes decreased heritability because more individuals survived and grew to reproduce at larger size. This surplus of adult individuals reproducing at large phenotype values, and producing relatively large offspring, increased the mean offspring phenotype and decreased the variance among offspring. This meant that potential parents were larger and more similar at birth (Table 1). As a consequence, the variance in parent birth phenotype decreased. However, the covariance between parent and offspring birth phenotype decreased substantially. Due to the overall higher survival probabilities and the reduced viability selection, more parents with small birth phenotypes grew to large adult sizes where they gave birth to large offspring. Defined by the ratio of this covariance and the parent birth phenotypic variance, heritability overall decreased for both species as the survival slope increased (Table 1).

Increasing the survival slopes for Soay sheep lambs and yearlings had comparable effects (Table 1). However, perturbing the survival slope for roe deer yearlings had little effect because the decrease in the covariance between parent and offspring birth phenotype was relatively weak.

### Elasticities of heritability with respect to fertility selection

Changing fertility and fertility selection had much larger effects on heritability for Soay sheep than for roe deer (Figs. 2 and 3). For Soay sheep, fertility selection mostly operated through the probability of recruitment because the twinning rates were very low (Fig. 4). Another idiosyncrasy of the Soay sheep model was that the probability of recruitment for adults was similar across all phenotypes (Fig. 4). Therefore, fertility selection in the adult age-class was low. Introducing fertility selection by increasing the slope of this function by an infinitesimal change of about 0.01 had the largest effect on heritability of all perturbations for Soay sheep (Fig. 2). It decreased heritability by approximately 5 %.

Similarly to the effect of increasing the adult survival slope, increasing fertility and positive fertility selection among adult Soay sheep resulted in more adults reproducing at large phenotype values. This increased the mean of the stable cohort birth phenotype distribution and decreased its variance (Table 1). The increase in fertility selection resulted in an increase in the mean parent birth phenotype, but the overall increase in fertility also led to smaller adults having higher fertility and therefore to an increase in the variance in birth phenotype among parents. The offspring birth phenotype increased due to the increase in fertility selection, and the variance in offspring birth phenotype decreased. The parent and offspring birth phenotypic covariance therefore decreased substantially, which resulted in a large decrease in heritability (Table 1). In Soay sheep, lambs have a small probability of recruitment (Fig. 4). Increasing this probability, and simultaneously increasing fertility selection by increasing the slope of the function, increased heritability (Fig. 2). The increase in the slope drastically increased the parent-offspring birth phenotypic covariance, because small-born parents were now more likely to give birth at small lamb sizes to small offspring. More lambs reproducing also caused a decrease in the mean and an increase in the variance of all three distributions: potential parents, parents, and offspring were on average smaller and less similar at birth. As a result, heritability increased (Table 1).

For roe deer, altering the probability of recruitment had less pronounced effects (Fig. 3). Increasing fertility and decreasing fertility selection for adults, due to the linearisation effect (Fig. S3), increased heritability. Supposedly, the decrease in fertility selection resulted in a decrease in the mean stable cohort birth phenotype because smaller adults had an increase in the probability of recruitment and a higher increase in the probability of recruitment than larger adults. At the same time, the variance in the stable cohort birth phenotype decreased; potential parents were smaller and more similar when compared to the unperturbed model. This smaller mean and variance translated into smaller means and variances for both the parent and offspring birth phenotype. The parent-offspring covariance increased, because parents that were born small, and therefore took longer to grow to the asymptotic size than large-born parents, had a higher probability of recruitment as small adults, resulting in small-born parents having a higher probability to give birth to small offspring (Table 1).

### Elasticities of heritability with respect to growth

Changing both intercepts or slopes of the functions that determined the variances in growth had little effect on heritability for both species (Figs. 2 and 3). These perturbations changed the variances in parent and offspring birth phenotype and in the parent-offspring covariance to similar proportions, which then overall had no effect on heritability (Tables S1 and S2). These findings even held for Soay sheep, where the growth variances have slopes that deviate from 0 (Fig. 4).

Increasing the slope of the mean growth function for adult and senescent Soay sheep had pronounced negative effects on heritability (Fig. 2). Increasing the slope of the mean growth function in an IPM has generally two consequences. First, all individuals of the relevant age-class attain faster their asymptotic body mass, with this acceleration being even larger for large individuals. Second, the mean maximum body mass increases because the mean growth rate function crosses the y = x line at higher phenotype values.

When the adult mean growth slope for Soay sheep was increased, individuals grew faster to larger phenotype values at which they had higher survival. This caused an increase in the number of adults reproducing at large phenotype values and therefore increased the mean size of potential parents, but also the variance in size among them (Table 1).

The increase in variance may have been caused by the increase in the asymptotic size of adults, which increased the range of phenotype values over which adults reproduce. This change in the phenotype mean and variance of the potential parent cohort cascaded through and led to similar increases in the mean and variance of parent and offspring birth phenotypes. The increase in the asymptotic size of adults, and the increase in how fast individuals grow to this size, increased the number of individuals that reproduce at large phenotype values regardless of their birth phenotype. Consequently, the covariance between parent and birth phenotype decreased, which divided by a much larger variance in birth phenotype, resulted in lower heritability (Table 1).

In order to understand how increasing the mean growth slope for senescent Soay sheep influenced heritability, it is important to notice that the slope for mean inheritance is negative for senescent individuals; the larger they are, the smaller is their offspring (Fig. 4). As a result, shifting the asymptotic size for senescent individuals towards larger phenotype values, at which individuals had higher probabilities of recruitment and survival yet gave birth to smaller offspring, decreased the mean and the variance in the stable cohort birth phenotype distribution. The variance further decreased because the variance of the growth function decreases with body mass in the senescent age-class. The decrease in the mean and variance in the stable cohort birth phenotype distribution was again followed by similar developments in the parent and offspring birth phenotype distributions. However, overall heritability decreased (Table 1) because the parent-offspring covariance decreased even more.

For roe deer, only increasing the growth mean slope for yearlings had a notable effect on heritability, since most of the growth towards the asymptotic body mass occurs in this age class (Fig. 3). Increasing the slope of yearling growth resulted in individuals growing faster to larger sizes at which they recruited to the adult age-class. They therefore experienced higher probabilities of survival, recruitment, and twinning rates during the first time step as adults. Due to the positive slope of the mean inheritance function (Fig. 5), this increased the mean and decreased the variance in the stable cohort birth phenotype distribution. Since potential parents were larger and more similar at birth, plus they were exposed to higher growth rates during the yearling age-class, both parents and offspring were also larger and more similar in birth phenotype, which increased the parent-offspring covariance (Table 1). Divided by a smaller parental variance, this led to higher heritability.

### Elasticities of heritability with respect to parent-offspring phenotype inheritance

The only direct relationship between offspring and parent phenotype (at birth of offspring) is modelled by the inheritance function. Nevertheless, perturbing this relationship by increasing the slopes of the mean inheritance functions had only moderate effects on heritability (Figs. 2 and 3). Increasing the slope of the mean inheritance function for Soay sheep yearlings and roe deer adults increased the mean stable cohort birth phenotype and decreased the associated variance. It supposedly decreased the variance because both age-classes had high probabilities of recruitment for the upper part of the stable phenotype distribution of the respective age-class (Figs. 4 and 5). Offspring produced were therefore larger and more similar. Since potential parents started out larger and more similar, the mean birth phenotype of parents was also larger and its variance smaller. Since parents were larger, they gave birth to larger and more similar offspring. The covariance between parent and offspring phenotype increased, while the variance in parent birth phenotype decreased, resulting in higher heritabilities caused by these perturbations.

However, increasing the slope of the mean inheritance function for adult Soay sheep actually decreased heritability (Fig. 2) because it increased the variance of the parent birth distribution. Since both small and large adult Soay sheep have about the same number of offspring due to the probability of recruitment being almost constant, increasing the slope of the mean inheritance function shifted the mean of the stable cohort birth phenotype distribution to larger sizes while spreading out the distribution. Since the stable cohort was larger and less similar in birth phenotype, the mean and variance in the parent birth phenotype distribution also increased. The covariance between parent and offspring birth phenotype also increased, but less than the variance in parent birth phenotype, which overall resulted in a lower heritability.

## Discussion

By studying two species of large vertebrates in an IPM framework, we have demonstrated how the effect of all four processes – viability and fertility selection, growth, and inheritance – on biometric heritability can be quantified and compared. Our results show that viability and fertility selection influence heritability more than growth and inheritance, which dampen or amplify the effect of selection. Interestingly, inheritance played the least important role. Our method allows us to understand if these processes influence heritability by influencing the phenotypic covariance among offspring and parents, the variance among parents, or both. Generally, processes that lead to individuals giving birth to offspring of different sizes decrease heritability. Accordingly, we estimated heritability of body mass at first census age to be lower in Soay sheep than in roe deer. Within-individual variation in offspring body mass is 67% and 44% for Soay sheep and roe deer, respectively. While Soay sheep can reproduce at different sizes in different age classes, giving birth to offspring of different sizes, roe deer mostly reproduce at sizes close to the mean asymptotic body size, resulting in more similar offspring. In the following, we discuss our main findings and the use of IPMs to study heritability in free-living animal populations. First, however, we summarise some general insights that provide guidance in distilling these insights from our varied results.

First, perturbing intercepts had smaller effects on heritability than perturbing slopes, because perturbing intercepts mostly affects the means of phenotype distributions, while perturbing slopes affects the means and the variances of phenotype distributions. Second, perturbations of the same parameter, but for different age classes, do not always change processes in the same way. Increasing the slope of the survival function, for example, increased viability selection for Soay sheep lambs yet decreased it for adults. Indeed, the slope of the survival function is not directly a measure of the strength of the viability selection. Their particular effects depend on the shape of the functions and on the stable age-class phenotype distributions (see the result part on Elasticities of heritability with respect to viability selection and Fig. S3). Third, the different functions interact. An increase in viability selection, for example, increases mean offspring size depending on the slope of the mean inheritance function. Finally, the more the functions vary among the different age classes, the less predictable are their interactions. This may explain why we see more and mostly stronger responses in heritability to perturbations for Soay sheep when compared to roe deer (Figs. 4 and 5).

### Effects of viability and fertility selection

Decreasing viability selection generally decreased heritability, as we have observed for both Soay sheep and roe deer in the adult and senescent age classes. This is because more parents of different birth phenotypes survive to grow and give birth to large offspring. In reality, however, strong changes in viability selection acting on adult Soay sheep and roe deer are unlikely to occur (Gaillard et al., 2000). More realistic is variation in viability selection among young individuals such as Soay sheep lambs, which we have observed would result in a decrease in heritability. In both species, viability selection up until the census age (8 months in roe deer, 4 months in Soay sheep) is part of the fertility selection, because the parental probabilities of recruitment and twinning include the probability for offspring of surviving to the census age. Increasing fertility selection had the strongest heritability-decreasing effect for Soay sheep. Many studies found that heritability is generally lower under poor than under favourable environmental conditions (reviewed in Charmantier & Garant, 2005; Merilä & Sheldon, 2001). Wilson et al. (2006) showed that additive genetic variance decreased with increased viability selection on offspring phenotype in Soay sheep. In line with their findings, we observed that increasing adult and senescent fertility selection, which includes viability selection on offspring before first census, decreased heritability. Moreover, given positive viability selection among adults, increasing fertility selection among adults is also likely to decrease directly heritability because it reduces the probability that small individuals reproduce.

### Effects of growth and inheritance

The effects of viability and fertility selection on heritability are mediated by ontogeny. If individuals grow slowly, then any effect of selection on heritability has more time to act via the still-growing individuals. If however individuals grow fast to large sizes, then selection has less opportunity to influence heritability. Therefore, we observed a decrease in heritability with an increase in the mean growth rate of Soay sheep adults. Furthermore, we found that fertility selection had less effect on heritability of roe deer than Soay sheep because roe deer were recruited at 8 months of age when most growth had already taken place (~ 70% of the asymptotic mass, Hewison et al., 2011).

Inheritance captures processes that directly relate variation in parental body condition and other maternal effects to offspring birth phenotype. Despite it being the only direct relationship between parental and offspring phenotype, the effect of inheritance on heritability is small. The mean inheritance function determines the variation in offspring phenotypes, which then results in the variation among parents through growth and selection processes. We found that increasing the slope of the mean inheritance function always increased the parent-offspring covariance. Remarkably though, this did not always result in an increase in heritability. For Soay sheep adults, the increase in phenotypic variance among potential parents also resulted in an increase in the variance in parent birth phenotype, which overall decreased heritability. The variance around the mean inheritance function captures reproductive allocation. Changing this variance has little effect on heritability because it changes the variances in parent phenotype and the parent-offspring covariances to similar proportions. In some species such as wild boars, individuals allocate resources differently to siblings as an evolutionary bet hedging strategy, called coin-flipping (Kaplan & Cooper, 1984), to minimise the variance in reproductive success (Gamelon et al., 2013). Our results show that this has no effect on heritability in roe deer and Soay sheep, but may play a role in species with larger litter sizes.

### Evolutionary and demographic effects on heritability estimates

Estimates of heritability for roe deer and soay sheep were within the range of heritabilities estimated from animal model (Soay sheep: 0.20 here vs 0.03-0.32 from animal model,Wilson et al. 2006; roe deer: 0.34 here vs. 0.07 and 0.44 in two other populations of roe deer, Quemere et. al, unpublished data). Because our model includes a growth and an inheritance function that link an offspring’s phenotype at birth to its mother’s phenotype at the age of reproduction (Chevin, 2015; Janeiro et al., 2016), it allows us to understand how and how much selection and growth can influence the parent-offspring regression. Heritability can increase both because the covariance among parents and offspring increases or because the total phenotypic variance among parents decreases. Our results show that demography influences always both the numerator and the denominator of this ratio. As illustrated by our results, the evolution of a phenotypic continuous trait is a complex interaction between transmission and selection. The pool of potential future parents used to quantify the total phenotypic variance and to estimate heritability is directly influenced by viability and fertility selection (Hadfield, 2008). The influence of selection on heritability has already been recognized in quantitative genetics (Hadfield, 2008; Nakagawa & Freckleton, 2008; Steinsland et al., 2014). Heywood (2005) derived a theoretical decomposition of the evolution of a trait, taking selection into account. However, it remains difficult to quantify each component of the decomposition given the currently available data from natural systems. Given the large influence of selection in roe deer and Soay sheep, it might partly explain why the predictions of the breeder’s equation are more accurate in lab or domestic populations with controlled selection than in wild populations where many and variable selective pressures can influence a trait (Bonnet et al., 2017). Thus, IPMs present a relevant approach to study the evolution of a continuous trait in natural systems, taking into account the interaction between environment and the dynamics of the trait.

According to quantitative genetics, the parent-offspring phenotypic covariance represents exclusively genetic similarity because the genotype does not develop with age. While this definition can be used as measuring heritability for traits that remain fixed for life, it is questionable for traits that develop with age. Indeed, parental effects often influence offspring body mass or size in addition to genetic and environmental effects (Mousseau & Fox, 1998). Parental effects may change according to parental age, condition, and experience. As a result, offspring phenotype cannot be predicted from additive genetic and environmental effects only (Fig. 1). Under the assumption that body mass is determined by a genotype that remains constant throughout life plus some environmental variation, a regression slope of unity would be expected. However, the slope of the growth function, where body mass at age *a* is regressed against body mass at age *a* + 1, is often estimated to be about 0.7 in IPMs (Coulson et al., 2010; Plard et al., 2015). The two ways out of this conundrum are for body mass to be determined by different but correlated genetic effects at different ages (Chevin, 2015) or for developmental trajectories to be controlled by genetic effects. Research to date has revealed strong positive genetic covariances across ages (Wilson et al., 2005) suggesting that this high genetic correlation across ages is not an explanation for the low regression slopes observed when consecutive body masses are regressed against one another. Instead such traits are likely strongly determined by the environment. Because the same genotype can attain different body masses in different environments, genotype-by-environment processes are also likely to be influential in size-related traits. Then, if developmental trajectories are genetically determined a useful approach is to break transmission down into contributions from growth increments, survival, reproduction and the correlation between parents and offspring. To better understand the evolution of phenotypic traits, we need to focus on the heritability of trajectories over life. In the future, our findings hopefully will spark further advances into understanding evolution in natural systems by challenging empiricists, eco-evolutionary, and quantitative geneticists to join forces.

## Acknowledgements

JAB acknowledges funding from the International Max Planck Research Network on Aging (MaxNetAging). TC acknowledges support from the NERC and an ERC advanced grant. ST was partly supported by US NIA grant R24AG039345. We thank Jason Matthiopoulos, Ben Sheldon, Michael Morrissey for helpful comments.

## Supporting information

**Appendix S1: Methods A1–A9** Details of the analytical framework.

**Table S1** Change in key population parameters when model parameters are perturbed (Soay sheep).

**Table S2** Change in key population parameters when model parameters are perturbed (roe deer).

**Figure S1** Comparison of the analytically and numerically computed elasticities of heritability for Soay sheep.

**Figure S2** Comparison of the analytically and numerically computed elasticities of heritability for roe deer.

**Figure S3** Schematic of the linearisation effect.

**Code S1** R code for analytically computing heritability and its elasticities. Download from http://tinyurl.com/gr8fgju.

## Supporting Information

### Appendix S1: Methods

#### A1 Parent-offspring covariance

The expected birth phenotype *Y*_0_ of offspring born to parents of age *a* and phenotype *X_a_* can be described by fitting a model of the form

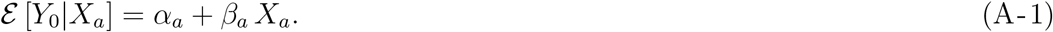

The expected offspring birth phenotype *Y*_0_ produced by a parent of age *a* and phenotype *z_i_* can also be calculated from the parent-offspring phenotype transition matrix as

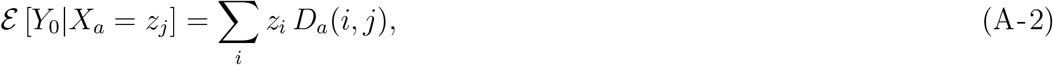

and so it follows that

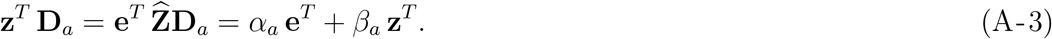

Since the lifetime production of offspring by a birth cohort is given by (8) in the main article, the probability that a newborn has a parent of age *a* is

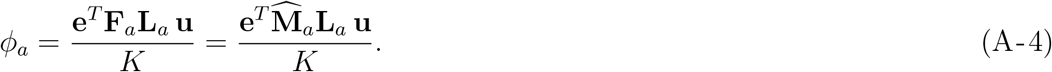

To derive a formula for the covariance between offspring birth phenotype *Y*_0_ and parent birth phenotype *X*_0_ that is computable in terms of **F**_*a,*_ **L**_*a*_, and **u**, we first derive the same covariance in terms of *ϕ_a_, α_a_*, and *β_a_*. Consider parents of age *a* and use (A-1) to see that

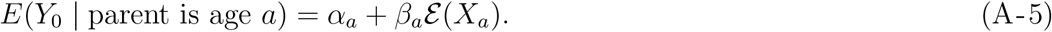

Hence, for lifetime reproduction, writing *α*̅ = Σ*_a_ ϕ_a_α_a_*,

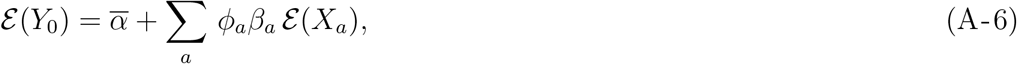

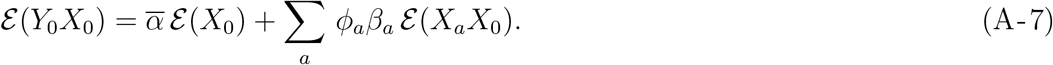

Using (A-6) and (A-7) the formula for the offspring-parent phenotype covariance in terms of the linear relationship in (A-1) is

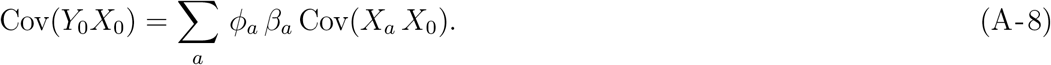

Next, to make the covariance computable in terms of our phenotype-demography matrices, the joint distribution of parent phenotype *X_a_* at age *a* and parent birth phenotype *X*_0_ is

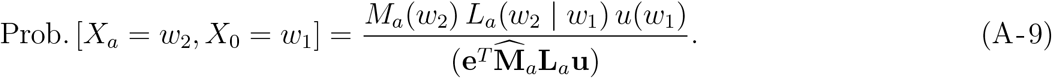

So given parents of age *a* with phenotype *X_a_* and birth phenotype *X*_0_,

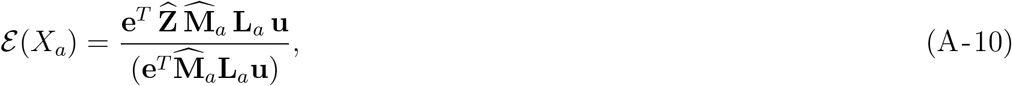

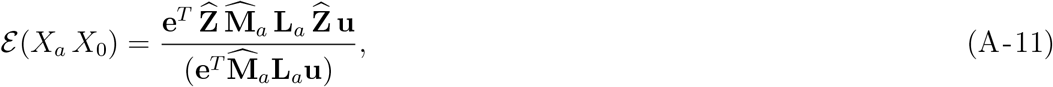

The joint distribution of offspring birth phenotype, parental phenotype at age *a*, and parent birth phenotype is

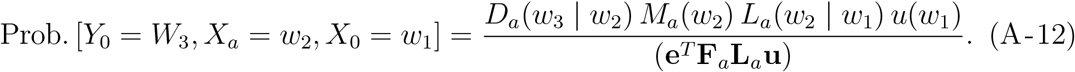

So we have

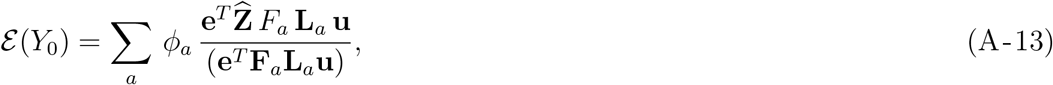

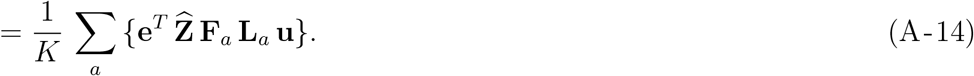

Next we have

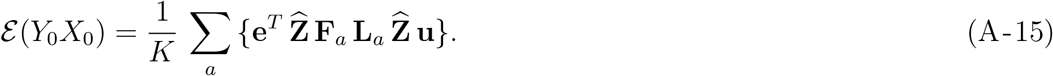

Now we use (A-14), (A-15), and (A-21) to obtain (10) in the main article, which is a formula for the parent-offspring birth phenotype covariance that is computable in terms of the phenotype-demography matrices.

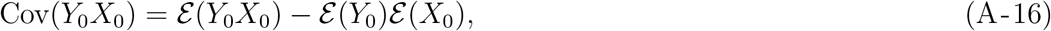

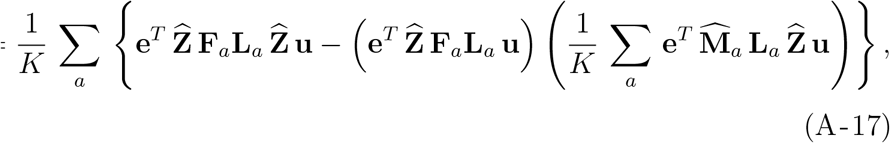

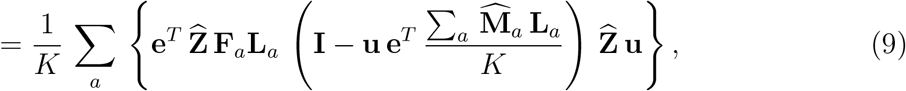

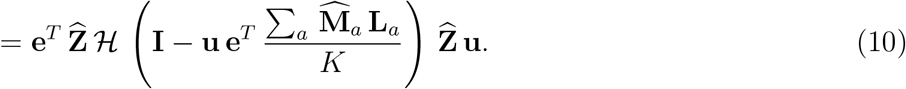

#### A2 Parent birth phenotype variance

In computing the variance, we need to account for the dispersion of lifetime reproduction across different ages. The parent cohort is born with the stable cohort birth phenotype distribution **u**. From (A-9) it follows that for parents aged *a* their average birth phenotype is

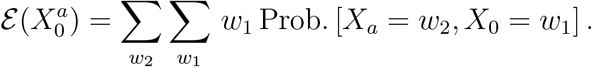

Therefore,

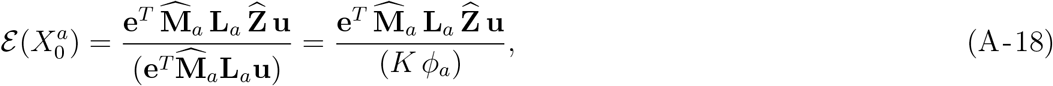

where the last equality uses (8) in the main article. The average squared birth phenotype for parents aged *a* is

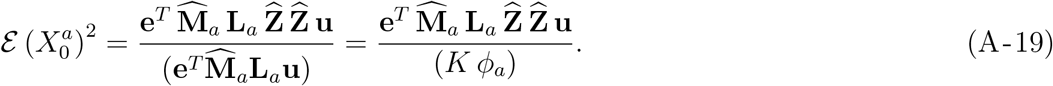

The variance of birth phenotype for parents of age *a* is

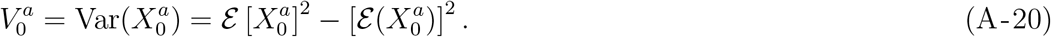

Now the average birth phenotype for all parents (i.e. of all ages) is

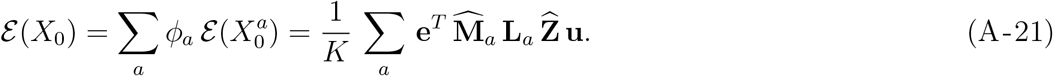

Finally, the variance of birth phenotype for all parents is

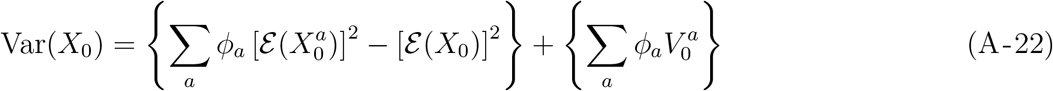

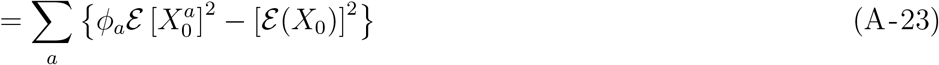

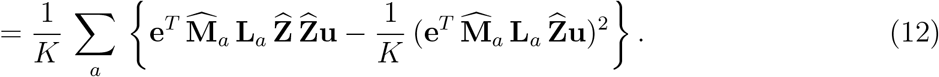

#### A3 Perturbation of u

For the unperturbed matrix **A**(*r*) we have defined the right eigenvector **u** as in (4) in the main article but also need the left eigenvector **v**,

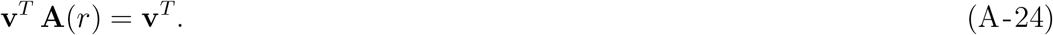

For convenience we normalize so that (**v***^T^* **u**) = 1. Then define the projection matrix

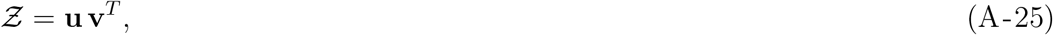

and the matrix

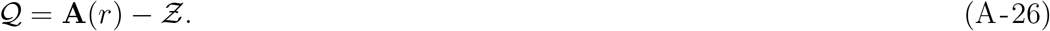

Now suppose that we change parameters with a small 0 *<* ∊ ≪ 1 so that

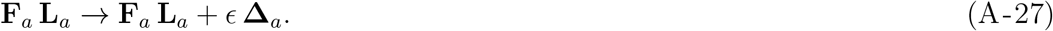

Then we have the resulting changes

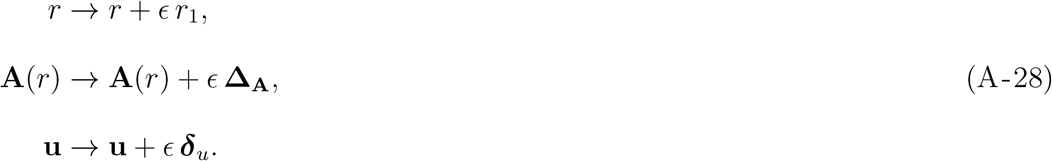

Note that the change ***δ***_*u*_ must be orthogonal to **u**, so that *Z****δ***_*u*_ = 0. From (4) in the main article **∆**_*A*_ and ***δ***_*u*_ are given by

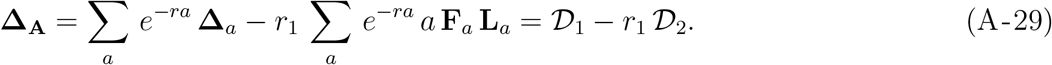

Clearly

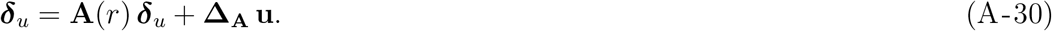

A standard argument (we multiply above on left by **v**^*T*^) yields the change

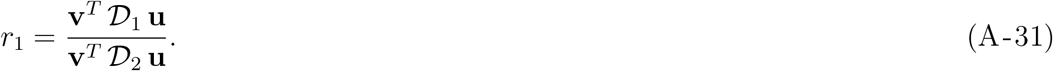

Next we observe that

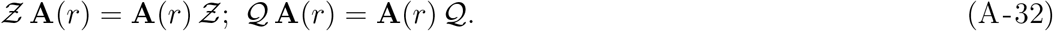

Recall that *Z****δ***_*u*_ = 0 so ***δ***_*u*_ = (**I** *− Z*) ***δ***_*u*_. This is why we multiply (A-30) on the left by (**I** *− Z*) to find

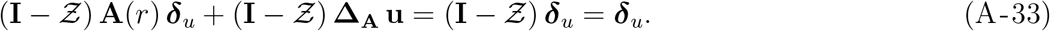

Next,

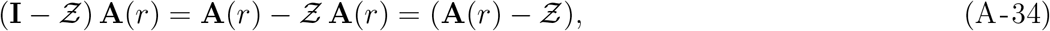

where we use (A-25). Hence (A-31) becomes

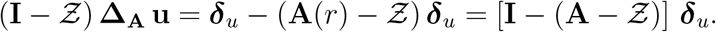

We now multiply across by [**I** *−* (**A** *− Z*)]^*−*1^ on both sides to obtain

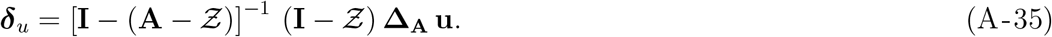

#### A4 Perturbation matrices

The perturbations in (A-27) come from changes in the fertility or survival matrices, as we explain in this subsection.

##### Selection on modifiers of fertility

A small change in fertility results from a change in total recruitment and/or in the phenotype distribution of offspring: thus at a given age *a*, there is a change in the fertility matrix **F**_*a*_ to (**F**_*a*_ + ∊ **∆_F_***a*), with

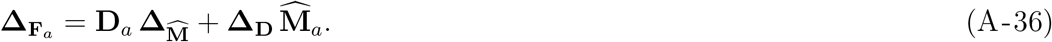

##### Selection on modifiers of phenotype transition rates

Phenotype transition rates **P**_*a*_ are made up of age-phenotype specific survival rates (*S_a_*(*x*)) and probabilities of growth from phenotype x’ to x (*G_a_*(*x|x^’^*)). A small change in one or more of these rates at age *a* means that we change **P**_*a*_ to **P***_a_ →* **P**_*a*_ + ∊ **∆_P_***a* with

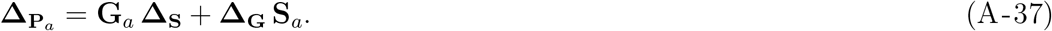

But notice that this change at age *a* in **P**_*a*_ will also change every survivorship **L**_*b*_ at ages *b > a*. The change in survivorship is and

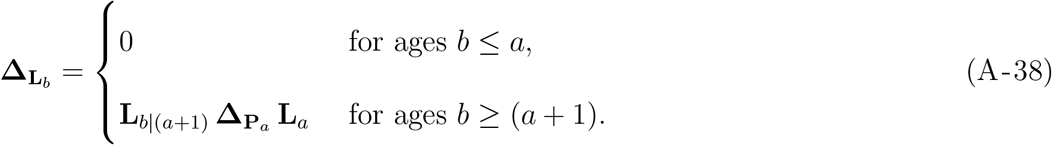

#### A5 Perturbation of parental variance

A small change in the phenotype-demography matrices as described in (A-27) results in a change in in the covariance between offspring birth phenotype and parent birth phenotype. Similarly, the change in the parent birth phenotype variance is

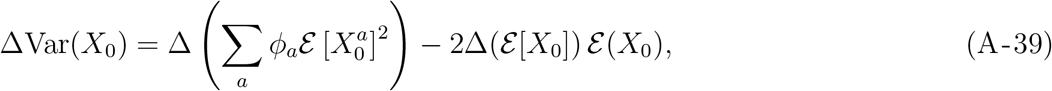

which gives with (A-19),

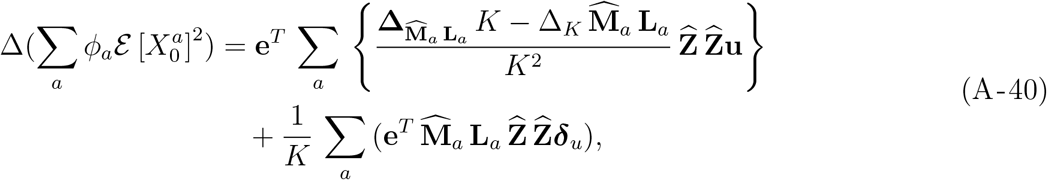

and

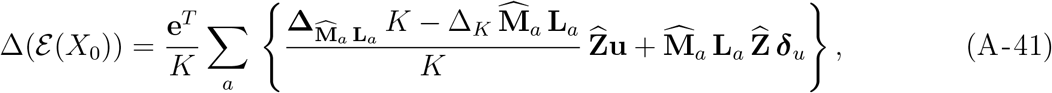

the equation (17) of the main article.

#### A6 Closure of terms

A general result we will use is: given a matrix 𝒬 and a real variable *w*,

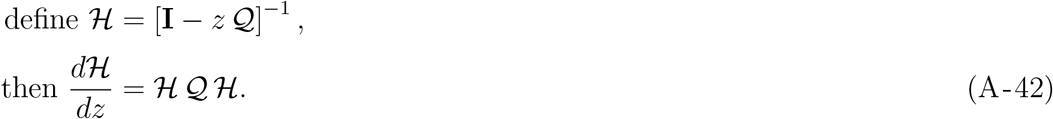

For our calculations in terms of the phenotype-demography matrices, suppose there is an age *m* after which fertility **F***_a_* = **F***_m_* and survival **P**_*a*_ = **P**_*m*_. Each sum in (A-29) can be written in two parts,

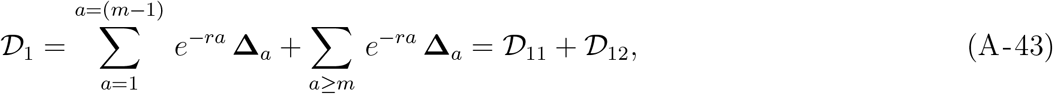

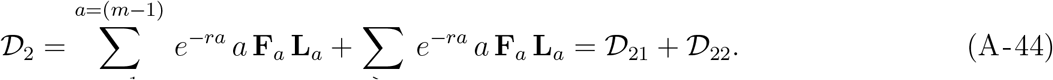

We have to do the sums to age (*m −* 1) as written. We seek explicit expressions for the sums that start at *m*. In the latter we have constant fertility and survival so

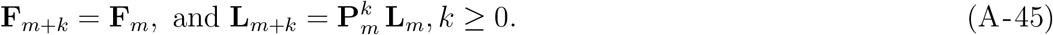

#### A7 Explicit form of 𝒟_22_

First we do the easier term 𝒟_22_, shown in (A-44). For a real variable *w*, define the matrix sum

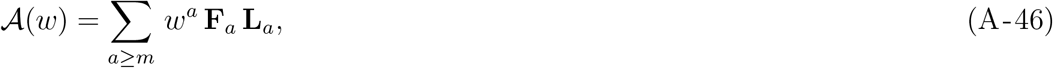

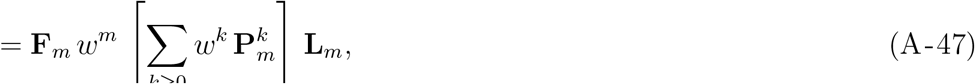

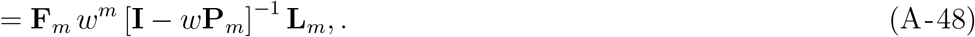

We can write *𝒜*(*w*) in a computable closed form in two steps:

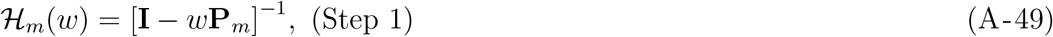

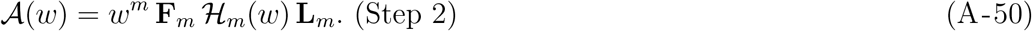

Next we use (A-46) to see that

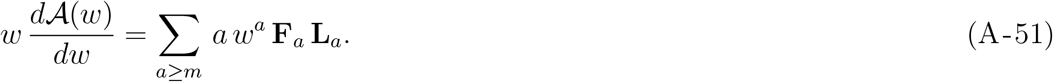

From (A-42) and (A-49) we find that

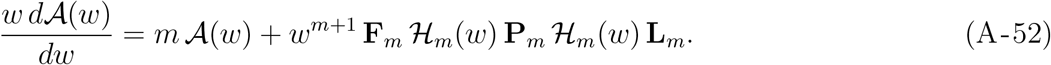

Hence the final expression: we use (A-49) and (A-50), and set *w* = *e^−r^*, to get

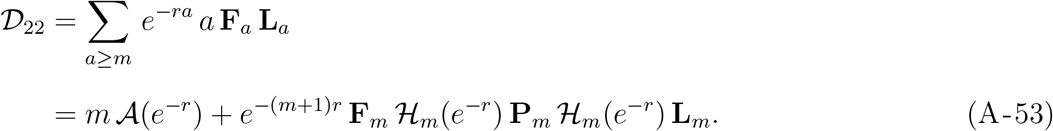

#### A8 Explicit form of 𝒟_12_

Second we turn to the more involved term 𝒟_12_ in (A-43) which is

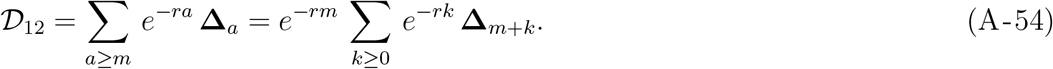

To sort this out we need to get explicit about perturbations to fertility and survival. In general we must consider the following perturbations:

> fertility **F**_*m*_ at ages *≥ m* changes to (**F**_*m*_ + ∊**∆**_*F*_);
>
> survival **P**_*m*_ at ages *≥ m* changes to (**P**_*m*_ + ∊ **∆**_*P*_);
>
> and cumulative survival **L**_*m*_ up to age *m* changes to (**L**_*m*_ + ∊ **∆**_*L*_). It is important to recognise that the change **∆**_*L*_ depends only on changes at ages *< m*.

Step 1: we need the change in 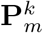. This is just the coefficient of *y* in (**P**_*m*_ + *y* **∆**_*P*_)*^k^* where *y* is just a real variable. For the present, we write this coefficient as **∆**_*Pk*_. Step 2: we recall (A-45): for *k ≥* 0 we have 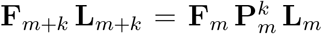 and therefore the linear change in this product is

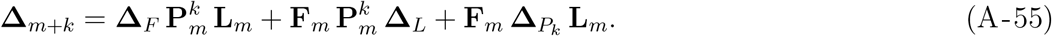

Step 3: we use (A-55) to rewrite the sum in (A-54) as

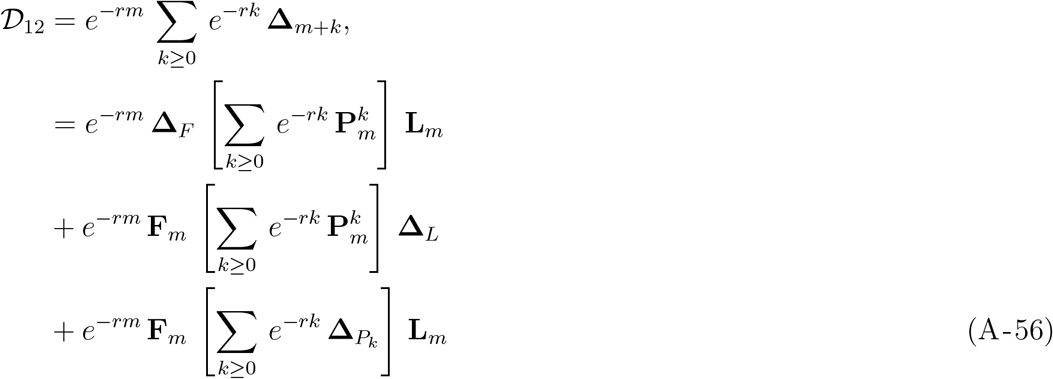

Step 4: we recall from (A-49) that ℋ_*m*_(*w*) = [**I** *− w***P**_*m*_]^*−*1^ and rewrite (A-56) as

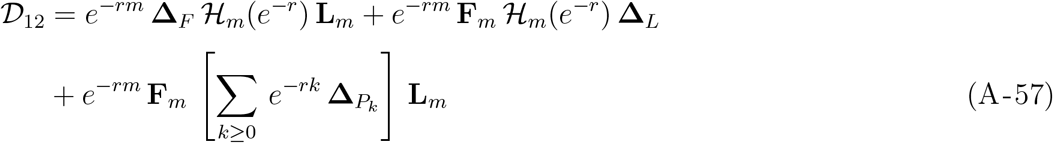

Step 5: now we just want a closed form for the sum in the second line of (A-57). Recall that **∆**_*Pk*_ is just the coefficient of *y* in (**P**_*m*_ + *y* **∆**_*P*_)^*k*^. So we define the function

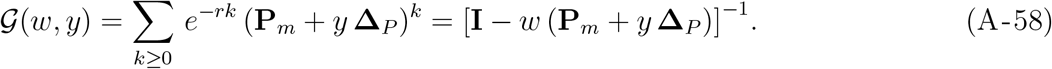

Then we must have

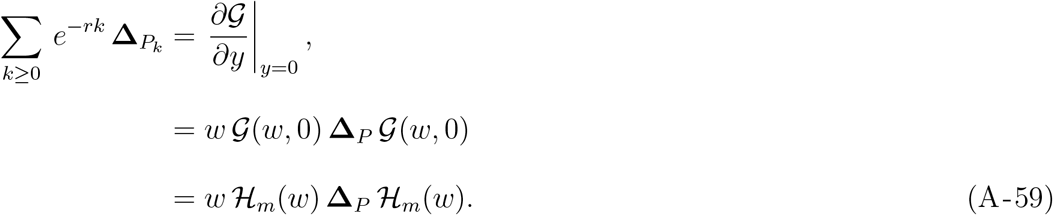

Finally, using (A-59) in (A-57) we have

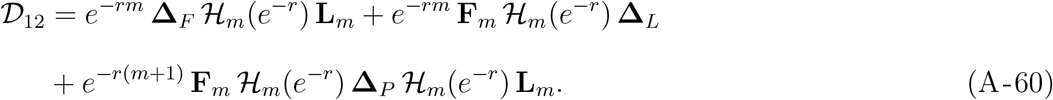

For the final expressions, we use (A-43) and (A-44). For the sums up to age (*m −* 1) we use 𝒟_11_, 𝒟_21_ as written. Then we use (A-60) to get 𝒟_12_ and (A-53) to get 𝒟_12_.

#### A9 Perturbation of transition densities

Growth and parent-offspring transition involve transition densities for a variable *y* conditional on a known variable *x*, usually taken to be proportional to a standard normal *f* with a mean *μ*(*x*) and variance *s*(*x*). The values of *y* are constrained to an interval [*A, B*] and we call the constrained transition density *f*̂,

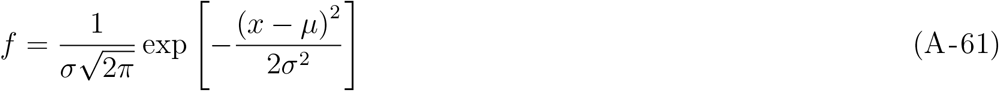

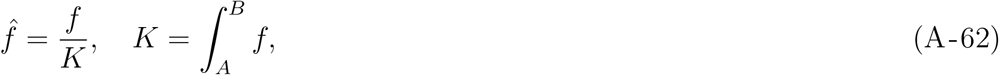

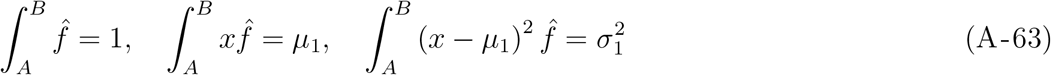

The mean *μ* (*x*) and variance *s*(*x*) depend on parameters that we call *θ*. A change in any parameter *θ* affects *f*̂ via *μ* or *s*, so here we simply consider changes in the latter.

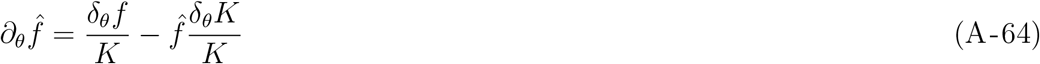

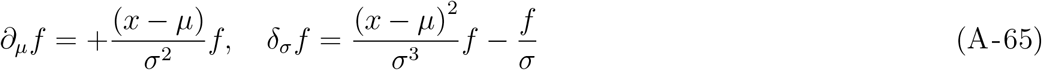

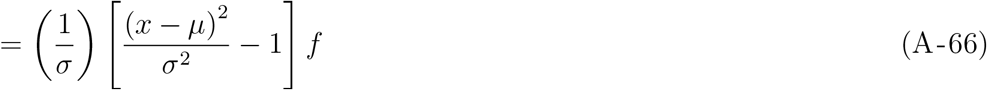

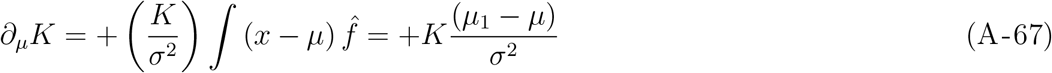

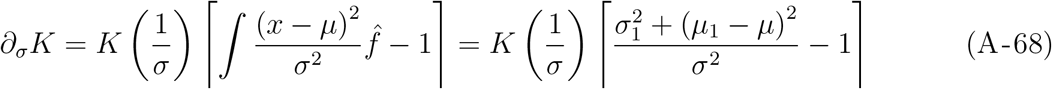

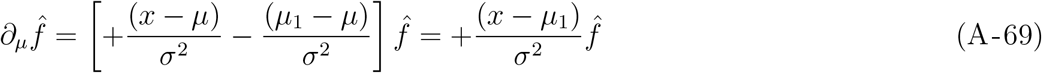

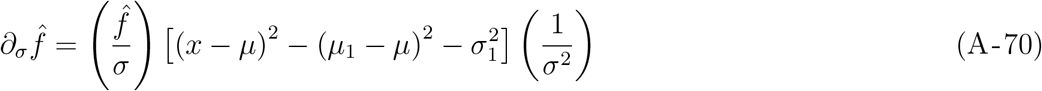

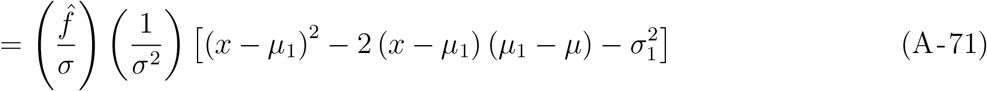

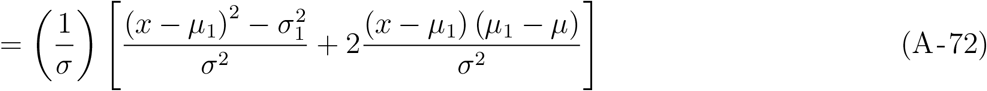

The final equations may be used to construct the perturbation matrices in (A-27)/(13).

### Appendix: Supplemental tables

**Table S1:** Change in key population parameters (%): expected and variance values of the stable cohort phenotype distribution (*X_c_*_0_), the stable parent (at own) birth phenotype distribution (*X*_0_) and offspring birth phenotype distribution (*Y*_0_) when selected model parameters are perturbed (Soay sheep). Note that shown values were calculated numerically by adding 0.01 to model parameters.

| | $\mathcal{E}(X_{c_0})$ | $\text{Var}(X_{c_0})$ | $\mathcal{E}(X_0)$ | $\text{Var}(X_0)$ | $\mathcal{E}(Y_0)$ | $\text{Var}(Y_0)$ | $\text{Cov}(Y_0X_0)$ | $h^2$ |
| --- | --- | --- | --- | --- | --- | --- | --- | --- |
| SurvIntL* | 0.00 | -0.03 | -0.02 | 0.00 | 0.01 | -0.11 | -0.46 | -0.46 |
| SurvIntY | 0.00 | -0.01 | -0.01 | 0.01 | 0.00 | -0.03 | -0.13 | -0.14 |
| SurvIntA | 0.00 | -0.01 | 0.00 | 0.00 | 0.01 | -0.05 | -0.21 | -0.20 |
| SurvIntS | 0.00 | -0.01 | 0.00 | -0.01 | 0.00 | -0.03 | -0.11 | -0.10 |
| SurvSlpL | 0.00 | -0.30 | -0.15 | -0.71 | 0.15 | -1.24 | -4.38 | -3.69 |
| SurvSlpY | 0.01 | -0.09 | -0.07 | 0.05 | 0.05 | -0.38 | -1.73 | -1.79 |
| SurvSlpA | 0.04 | -0.14 | -0.01 | -0.11 | 0.15 | -0.94 | -3.75 | -3.65 |
| SurvSlpS | -0.02 | -0.27 | -0.03 | -0.32 | 0.04 | -0.83 | -2.67 | -2.36 |
| TwIntL | 0.00 | 0.00 | 0.00 | 0.00 | 0.00 | 0.00 | 0.00 | 0.00 |
| TwIntY | 0.00 | 0.00 | 0.00 | 0.00 | 0.00 | 0.00 | 0.00 | 0.00 |
| TwIntA | 0.00 | 0.00 | 0.00 | 0.00 | 0.00 | 0.00 | 0.00 | 0.00 |
| TwIntS | 0.00 | 0.00 | 0.00 | 0.00 | 0.00 | 0.00 | 0.00 | 0.00 |
| TwSlpL | 0.00 | 0.00 | 0.00 | 0.00 | 0.00 | 0.00 | 0.00 | 0.00 |
| TwSlpY | 0.00 | 0.00 | 0.00 | 0.00 | 0.00 | 0.00 | 0.00 | 0.00 |
| TwSlpA | 0.00 | 0.00 | 0.00 | 0.00 | 0.00 | -0.01 | -0.01 | -0.01 |
| TwSlpS | 0.00 | -0.01 | 0.00 | -0.01 | 0.00 | -0.04 | -0.13 | -0.11 |
| PrIntL | -0.03 | 0.24 | -0.01 | 0.11 | -0.03 | 0.22 | 0.66 | 0.54 |
| PrIntY | 0.00 | -0.01 | 0.00 | 0.00 | 0.00 | -0.02 | 0.04 | 0.04 |
| PrIntA | 0.01 | -0.01 | 0.00 | 0.00 | 0.02 | -0.08 | -0.22 | -0.22 |
| PrIntS | 0.00 | -0.02 | 0.00 | -0.02 | 0.00 | -0.05 | -0.15 | -0.13 |
| PrSlpL | -0.41 | 2.84 | -0.13 | 1.37 | -0.35 | 2.59 | 4.53 | 3.12 |
| PrSlpY | -0.08 | -0.20 | -0.04 | -0.10 | -0.04 | -0.34 | 0.53 | 0.62 |
| PrSlpA | 0.15 | -0.03 | 0.14 | 0.11 | 0.40 | -1.58 | -4.11 | -4.21 |
| PrSlpS | -0.04 | -0.41 | -0.05 | -0.46 | 0.04 | -1.13 | -3.76 | -3.31 |
| GrMIntL | 0.01 | 0.01 | 0.00 | 0.02 | 0.01 | 0.00 | -0.01 | -0.03 |
| GrMIntY | 0.01 | 0.02 | 0.01 | 0.03 | 0.01 | 0.02 | -0.01 | -0.03 |
| GrMIntA | 0.01 | 0.07 | 0.02 | 0.09 | 0.02 | 0.04 | -0.06 | -0.15 |
| GrMIntS | 0.00 | -0.01 | 0.00 | -0.01 | 0.00 | -0.02 | -0.08 | -0.07 |
| GrMSlpL | 0.07 | 0.19 | 0.08 | 0.17 | 0.09 | 0.02 | 0.67 | 0.49 |
| GrMSlpY | 0.10 | 0.43 | 0.12 | 0.45 | 0.11 | 0.30 | 0.40 | -0.05 |
| GrMSlpA | 0.30 | 1.55 | 0.39 | 2.01 | 0.38 | 0.90 | -0.71 | -2.67 |
| GrMSlpS | -0.04 | -0.15 | -0.03 | -0.14 | -0.01 | -0.57 | -2.09 | -1.95 |

Table S1: continued.
| | $\mathcal{E}(X_{c_0})$ | $\text{Var}(X_{c_0})$ | $\mathcal{E}(X_0)$ | $\text{Var}(X_0)$ | $\mathcal{E}(Y_0)$ | $\text{Var}(Y_0)$ | $\text{Cov}(Y_0 X_0)$ | $h^2$ |
| --- | --- | --- | --- | --- | --- | --- | --- | --- |
| GrVIntL | 0.00 | 0.00 | 0.00 | 0.00 | 0.00 | 0.00 | 0.00 | 0.00 |
| GrVIntY | 0.00 | 0.00 | 0.00 | 0.00 | 0.00 | 0.00 | 0.00 | 0.00 |
| GrVIntA | 0.00 | 0.00 | 0.00 | 0.01 | 0.00 | 0.00 | 0.01 | 0.00 |
| GrVIntS | 0.00 | 0.00 | 0.00 | 0.00 | 0.00 | 0.00 | 0.00 | 0.00 |
| GrVSlpL | 0.01 | 0.03 | 0.01 | 0.04 | 0.01 | 0.03 | 0.11 | 0.07 |
| GrVSlpY | 0.00 | 0.03 | 0.00 | 0.05 | 0.00 | 0.02 | 0.05 | -0.01 |
| GrVSlpA | 0.00 | 0.09 | 0.01 | 0.21 | 0.00 | 0.08 | 0.12 | -0.09 |
| GrVSlpS | 0.00 | 0.00 | 0.00 | 0.00 | 0.00 | -0.01 | -0.07 | -0.07 |
| InMIntL | 0.01 | -0.11 | 0.00 | -0.02 | 0.01 | -0.10 | -0.06 | -0.04 |
| InMIntY | 0.01 | -0.01 | 0.00 | -0.02 | 0.01 | -0.01 | 0.05 | 0.07 |
| InMIntA | 0.06 | 0.10 | 0.05 | 0.07 | 0.06 | 0.07 | -0.04 | -0.12 |
| InMIntS | 0.01 | 0.01 | 0.01 | -0.01 | 0.02 | 0.00 | 0.00 | 0.01 |
| InMSlpL | 0.09 | -1.35 | 0.00 | -0.30 | 0.08 | -1.25 | 0.02 | 0.33 |
| InMSlpY | 0.13 | -0.10 | 0.09 | -0.29 | 0.12 | -0.17 | 1.45 | 1.75 |
| InMSlpA | 1.30 | 2.55 | 1.25 | 2.13 | 1.35 | 1.87 | 1.52 | -0.60 |
| InMSlpS | 0.30 | 0.28 | 0.26 | -0.13 | 0.38 | 0.16 | 0.19 | 0.32 |
| InV1L | 0.00 | 0.01 | 0.00 | 0.00 | 0.00 | 0.01 | -0.01 | 0.00 |
| InVIntY | 0.00 | 0.01 | 0.00 | 0.01 | 0.00 | 0.01 | 0.01 | 0.00 |
| InVIntA | 0.00 | 0.10 | 0.01 | 0.13 | 0.00 | 0.10 | 0.12 | -0.01 |
| InVIntS | 0.00 | 0.02 | 0.00 | 0.02 | 0.00 | 0.03 | 0.02 | 0.00 |
| InVSlpL | 0.00 | 0.15 | 0.00 | -0.05 | 0.00 | 0.13 | -0.06 | -0.01 |
| InVSlpY | 0.00 | 0.22 | 0.02 | 0.17 | 0.00 | 0.20 | 0.18 | 0.01 |
| InVSlpA | 0.01 | 2.13 | 0.20 | 2.86 | 0.02 | 2.17 | 2.68 | -0.18 |
| InVSlpS | 0.00 | 0.50 | 0.05 | 0.55 | 0.00 | 0.62 | 0.53 | -0.02 |

\*Model parameters were perturbed upwards by adding 0.01. Quantities were calculated numerically. Model parameter names are written in CamelCase forming compounds of the following abbreviations in order of appearance: survival (Surv), intercept (int), lambs (L), yearlings (Y), prime-aged adults (A), senescents (S), slope (slp), twinning rate (Tw), probability of reproduction (Pr), growth (Gr), mean (M), variance (V), inheritance (In).

**Table S2:** Change in key population parameters (%): expected and variance values of the stable cohort phenotype distribution (*X_c_*_0_), the stable parent (at own) birth phenotype distribution (*X*_0_) and offspring birth phenotype distribution (*Y*_0_) when selected model parameters are perturbed (roe deer). Note that shown values were calculated numerically by adding 0.01 to model parameters.

| | $\mathcal{E}(X_0)$ | $\text{Var}(X_0)$ | $\mathcal{E}(X_0^P)$ | $\text{Var}(X_0^P)$ | $\mathcal{E}(Y)$ | $\text{Var}(Y)$ | $\text{Cov}(Y X_0^P)$ | $h^2$ |
| --- | --- | --- | --- | --- | --- | --- | --- | --- |
| SurvIntY* | -0.01 | 0.00 | -0.02 | 0.01 | 0.00 | 0.01 | 0.02 | 0.00 |
| SurvIntA | 0.00 | -0.01 | 0.00 | 0.00 | 0.02 | 0.00 | -0.30 | -0.30 |
| SurvIntS | 0.00 | 0.01 | 0.00 | 0.01 | 0.01 | 0.01 | -0.05 | -0.06 |
| SurvSlpY | -0.11 | 0.00 | -0.17 | -0.12 | -0.03 | 0.07 | -0.09 | 0.02 |
| SurvSlpA | 0.13 | -0.13 | 0.02 | -0.16 | 0.36 | -0.10 | -6.02 | -5.87 |
| SurvSlpS | 0.07 | 0.20 | 0.08 | 0.21 | 0.16 | 0.31 | -1.32 | -1.52 |
| TwIntY | 0.00 | 0.00 | 0.00 | 0.00 | 0.00 | 0.00 | 0.00 | 0.00 |
| TwIntA | -0.01 | -0.01 | -0.01 | -0.01 | 0.00 | 0.00 | 0.01 | 0.02 |
| TwIntS | 0.00 | 0.00 | 0.00 | 0.00 | 0.00 | 0.01 | -0.01 | -0.02 |
| TwSlpY | 0.00 | 0.00 | 0.00 | 0.00 | 0.00 | 0.00 | 0.00 | 0.00 |
| TwSlpA | -0.12 | -0.20 | -0.12 | -0.24 | -0.06 | -0.15 | -0.12 | 0.12 |
| TwSlpS | 0.04 | 0.12 | 0.04 | 0.12 | 0.06 | 0.15 | -0.33 | -0.45 |
| PrIntY | 0.00 | 0.00 | 0.00 | 0.00 | 0.00 | 0.00 | 0.00 | 0.00 |
| PrIntA | -0.02 | 0.00 | -0.02 | 0.00 | -0.01 | 0.01 | 0.08 | 0.08 |
| PrIntS | 0.00 | 0.00 | 0.00 | 0.00 | 0.00 | 0.01 | -0.02 | -0.03 |
| PrSlpY | 0.00 | 0.00 | 0.00 | 0.00 | 0.00 | 0.00 | 0.00 | 0.00 |
| PrSlpA | -0.27 | -0.14 | -0.29 | -0.15 | -0.16 | -0.03 | 0.86 | 1.01 |
| PrSlpS | 0.04 | 0.12 | 0.05 | 0.12 | 0.08 | 0.14 | -0.54 | -0.66 |
| GrMIntY | 0.02 | -0.05 | 0.02 | -0.05 | 0.02 | -0.05 | -0.07 | -0.02 |
| GrMIntA | 0.05 | 0.09 | 0.05 | 0.09 | 0.06 | 0.09 | -0.01 | -0.10 |
| GrMIntS | 0.00 | 0.02 | 0.00 | 0.02 | 0.00 | 0.03 | 0.01 | -0.01 |
| GrMSlpY | 0.32 | -0.68 | 0.28 | -0.69 | 0.30 | -0.72 | 0.50 | 1.21 |
| GrMSlpA | 1.25 | 2.83 | 1.26 | 2.72 | 1.52 | 2.81 | 2.33 | -0.38 |
| GrMSlpS | 0.07 | 0.44 | 0.08 | 0.50 | 0.13 | 0.80 | 0.31 | -0.19 |
| GrVIntY | 0.00 | 0.02 | 0.00 | 0.01 | 0.00 | 0.01 | 0.01 | 0.00 |
| GrVIntA | 0.00 | 0.05 | 0.00 | 0.05 | 0.00 | 0.05 | 0.05 | -0.01 |
| GrVIntS | 0.00 | 0.00 | 0.00 | 0.00 | 0.00 | 0.01 | 0.00 | 0.00 |
| GrVSlpY | 0.02 | 0.25 | 0.03 | 0.21 | 0.02 | 0.19 | 0.23 | 0.02 |
| GrVSlpA | 0.08 | 1.17 | 0.11 | 1.26 | 0.09 | 1.28 | 1.24 | -0.02 |
| GrVSlpS | 0.00 | 0.08 | 0.01 | 0.10 | 0.01 | 0.15 | 0.09 | 0.00 |
| InMIntY | 0.00 | 0.00 | 0.00 | 0.00 | 0.00 | 0.00 | 0.00 | 0.00 |
| InMIntA | 0.07 | -0.08 | 0.06 | -0.08 | 0.07 | -0.10 | -0.10 | -0.02 |
| InMIntS | 0.01 | 0.05 | 0.01 | 0.05 | 0.01 | 0.07 | 0.06 | 0.01 |
| InMSlpY | 0.00 | 0.00 | 0.00 | 0.00 | 0.00 | 0.00 | 0.00 | 0.00 |
| InMSlpA | 1.64 | -0.71 | 1.51 | -0.84 | 1.54 | -1.33 | 0.70 | 1.56 |
| InMSlpS | 0.16 | 1.28 | 0.19 | 1.50 | 0.25 | 1.95 | 1.67 | 0.17 |

| Table S2: continued. |  |  |  |  |  |  |  |  |
| --- | --- | --- | --- | --- | --- | --- | --- | --- |
| | $\mathcal{E}(X_0)$ | $\text{Var}(X_0)$ | $\mathcal{E}(X_0^P)$ | $\text{Var}(X_0^P)$ | $\mathcal{E}(Y)$ | $\text{Var}(Y)$ | $\text{Cov}(YX_0^P)$ | $h^2$ |
| InVIntY | 0.00 | 0.00 | 0.00 | 0.00 | 0.00 | 0.00 | 0.00 | 0.00 |
| InVIntA | 0.00 | 0.16 | 0.01 | 0.15 | 0.00 | 0.15 | 0.15 | 0.00 |
| InVIntS | 0.00 | 0.01 | 0.00 | 0.02 | 0.00 | 0.02 | 0.02 | 0.00 |
| InVSlpY | 0.00 | 0.00 | 0.00 | 0.00 | 0.00 | 0.00 | 0.00 | 0.00 |
| InVSlpA | 0.03 | 3.79 | 0.14 | 3.63 | 0.02 | 3.52 | 3.62 | -0.01 |
| InVSlpS | 0.00 | 0.35 | 0.01 | 0.50 | 0.00 | 0.59 | 0.49 | -0.01 |

\*Model parameters were perturbed upwards by adding 0.01. Quantities were calculated numerically. Model parameter names are written in CamelCase forming compounds of the following abbreviations in order of appearance: survival (Surv), intercept (Int), yearlings (Y), prime-aged adults (A), senescents (S), slope (slp), twinning rate (Tw), probability of reproduction (Pr), growth (Gr), mean (M), variance (V), inheritance (In).

### Appendix: Supplemental figures

**Figure S1:**
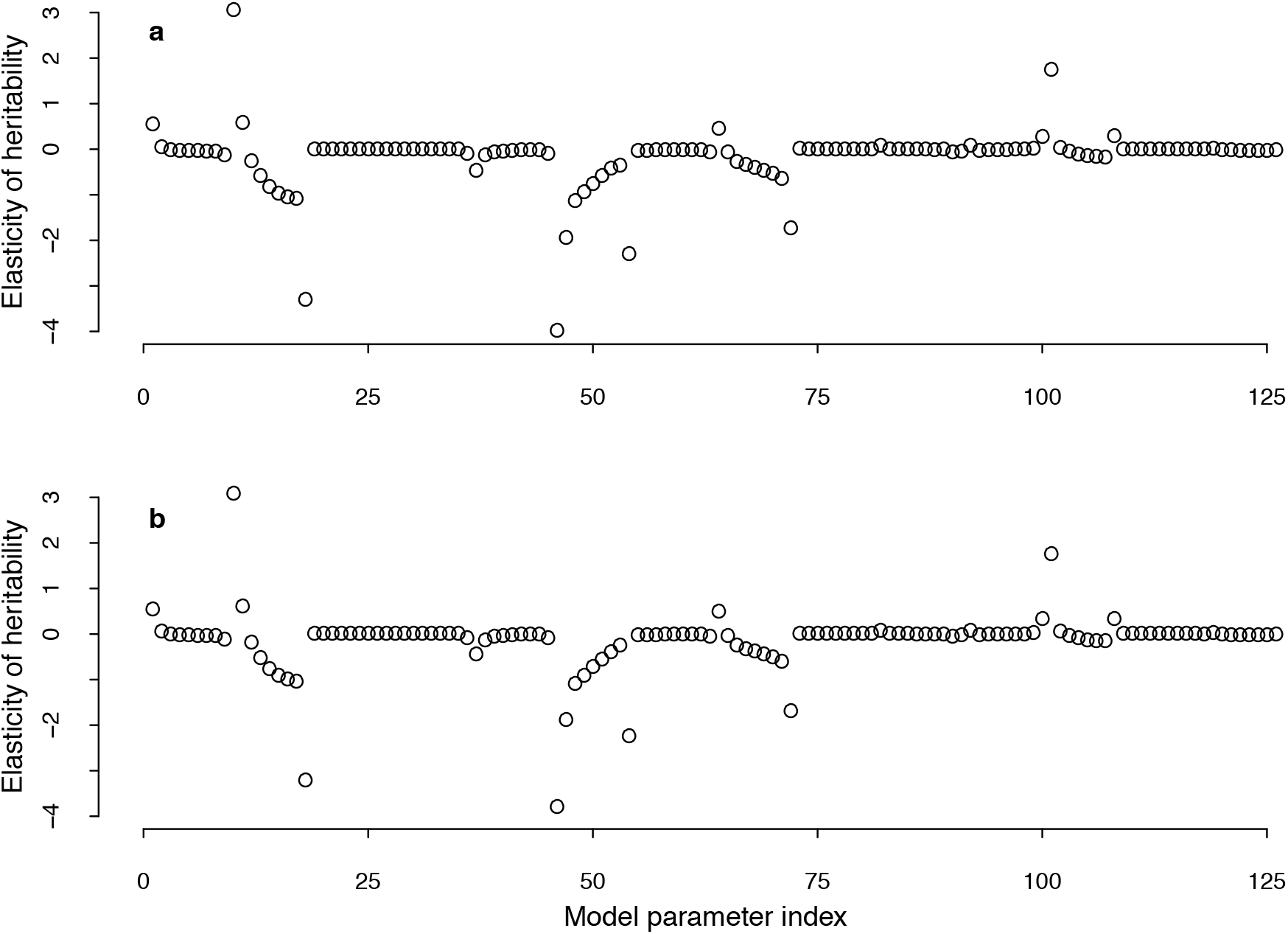
The analytically (a) and numerically (b) computed elasticities of heritability with respect to changes in the model parameters of the Soay sheep IPM. The two ways of computation give very similar results. Note that the values are not summed up within age classes.

**Figure S2:**
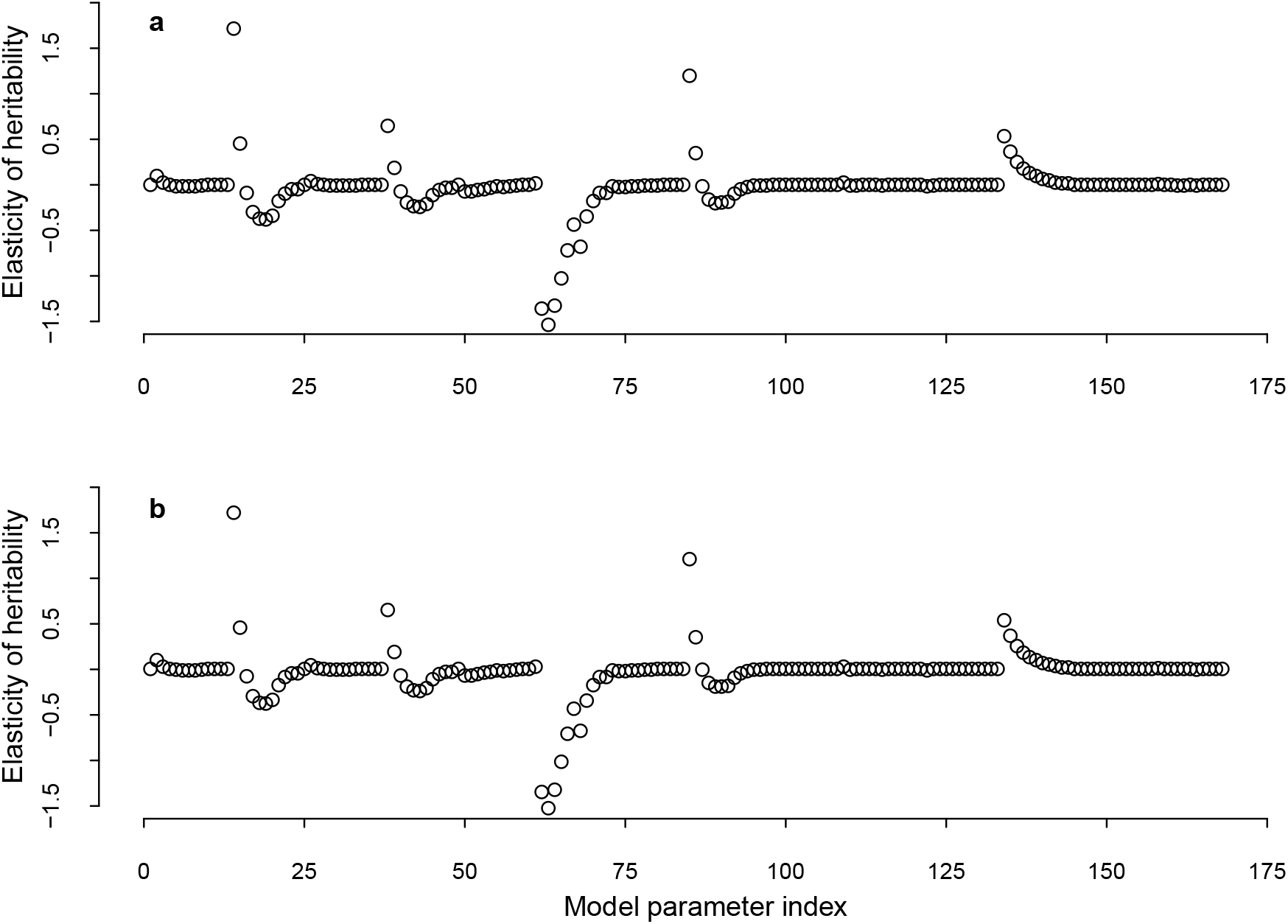
The analytically (a) and numerically (b) computed elasticities of heritability with respect to changes in the model parameters of the roe deer IPM. As for Soay sheep, the two ways of computation give very similar results.

**Figure S3:**
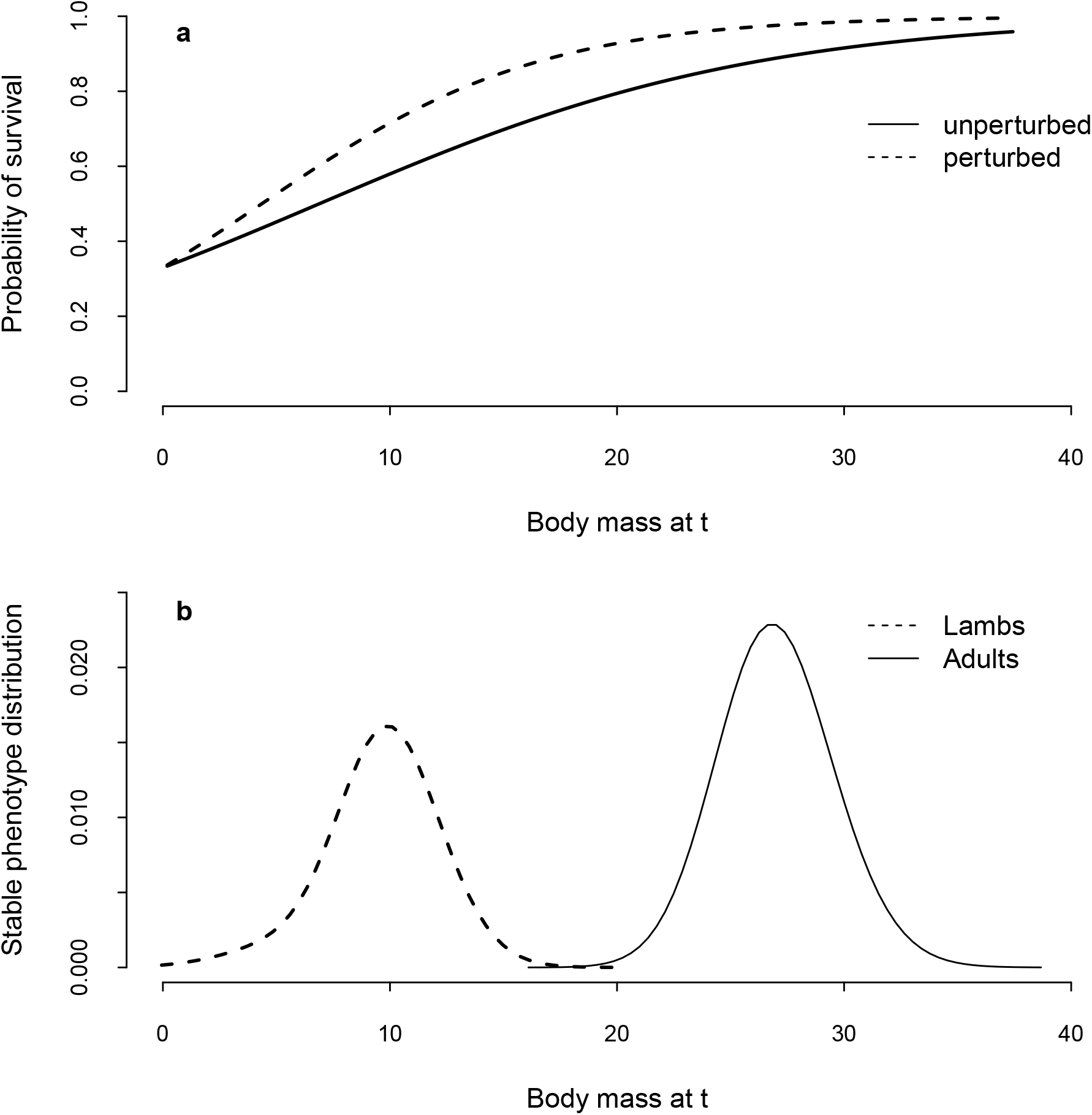
Schematic of the linearisation effect using a linearised survival function as an example. The effect of perturbing the slope of the survival function (a) on viability selection depends on the phenotype distribution (b). The same perturbation in (a) makes survival probabilities less equal across the lamb phenotype values and more equal across the adult phenotype values (b). Therefore the same perturbation increases viability selection for lambs and decreases viability selection for adults.

